# The landscape of S100B+ and HLA-DR+ dendritic cell subsets in tonsils at the single cell level via high-parameter mapping

**DOI:** 10.1101/369983

**Authors:** Maddalena Maria Bolognesi, Francesca Maria Bosisio, Marco Manzoni, Denis Schapiro, Riccardo Tagliabue, Mario Faretta, Carlo Parravicini, Ann M Haberman, Giorgio Cattoretti

## Abstract

Dendritic cells (DC) (classic, plasmacytoid, inflammatory) are an intense focus of interest because of their role in inflammation, autoimmunity, vaccination and cancer. We present a tissue-based classification of human DC subsets in tonsils with a high-parameter (>40 markers) immunofluorescent approach, cell type-specific image segmentation and the use of bioinformatics platforms. Through this deep phenotypic and spatial examination, classic cDC1, cDC2, pDC subsets have been further refined and a novel subset of DC co-expressing IRF4 and IRF8 identified. Based on unique tissue locations within the tonsil, and close interactions with T cells (cDC1) or B cells (cDC2), DC subsets can be further subdivided by correlative phenotypic changes associated with these interactions. In addition, monocytes and macrophages expressing HLA-DR or S100AB are identified and localized in the tissue. This study thus provides a whole tissue in situ catalog of human DC subsets and their cellular interactions within spatially defined niches.

## INTRODUCTION

Dendritic cells (DC) orchestrate a variety of adaptive immune responses by capturing and processing antigens for presentation to rare antigen-specific T cells (Banchereau and Steinman, 1998). Class II histocompatibility complex (HLA-DR) is crucial for effective antigen presentation and synapse formation with cognate T-cells and undergoes dynamic changes in subcellular localization and expression (Cella et al., 1997; Pierre et al., 1997). A master transactivator, CIITA, and transcription factors (TF) such as SPI1/PU.1, IRF8 and PRDM1 regulate HLA-DR in fully activated DC (Smith et al., 2011).

DCs respond themselves to the changes induced in the engaged T cells by releasing chemokines and cytokines, by modulating the repertoire of surface and cytoplasmic proteins such as CD40, CD80, CD86 (Banchereau and Steinman, 1998) and downregulate macrocytosis and newly acquired antigen processing, favoring the retention of peptide-charged HLA-DR on the surface (Wilson et al., 2004b). Engagement of innate immunity receptors (TLR7, −9) or chronic exposure to virus, downregulate specific TF in pDC, but not HLA-DR (Wu et al., 2008). Variegation of TF in mature cDC upon physiologic receptor or cell-cell engagement has not been reported.

DC are under TF control during commitment and differentiation from a common lympho-myeloid precursor (Merad et al., 2013). The Zbtb46 TF (Satpathy et al., 2012) and other TF (BCL11A, ID2, IRF4, IRF8, BCL6 (Guilliams et al., 2016; Ippolito et al., 2014; Zhang et al., 2014)) are required for lineage specification and maturation of the various subsets of DC (Belz and Nutt, 2012; Merad et al., 2013).

A consensus nomenclature divides mammalian DC into conventional dendritic cells type 1 (cDC1), type 2 (cDC2) and plasmacytoid DC (pDC), each with a distinctive set of prototypic biomarkers (CD141, CD1c, CD303)(Haniffa et al., 2012). and non-overlapping biological functions (Guilliams et al., 2016; Haniffa et al., 2012). Cells of monocyte-macrophage lineage, once admixed with DC because of the shared dendritic morphology, surface markers and some common functions, are now separately defined (Geissmann et al., 2010).

Initial phenotypic studies, validated and extended by single cell RNAseq (Villani et al., 2017) or high-parameter mass cytometry (Guilliams et al., 2016) have allowed for a further refinement in the subsetting of cDC (See et al., 2017; Villani et al., 2017). One of these studies has identified two members of a large protein family, S100A8 and A9, expressed in cDC2 (Villani et al., 2017). In the past another S100 proteins, S100B, has been used as tissue-based marker for cells with dendrites of various length and complexity (“dendritic” cells) which include interdigitating reticulum cells (IRC) (Takahashi et al., 1981), veiled cells, Langerhans cells of the epidermis (LC), and follicular dendritic cells (FDC).

S100B dendritic cells have been described interacting with radically different lymphoid cells: T-cells (Takahashi et al., 1998) and B-cell (Takahashi et al., 2006).

A tissue-based classification of human DC, including the S100B+ subset, is lacking, thus we embarked on an in situ analysis of S100B and HLA-DR positive cells in the normal lymphoid tonsil tissue, combining traditional immunofluorescence tissue staining with a powerful technique that enables a high-parameter characterization with a spatial computational analysis. This approach has empowered a detailed catalog of human DC subsets and their cellular interactions within spatially defined tissue niches.

## RESULTS

Antibody raised against bovine brain S100, preferentially recognizing S100B (henceforth named S100AB) decorate sparse cells of dendritic morphology, small lymphocytes, and follicular dendritic cells, as expected based on previous publications (Takahashi et al., 1981). >99% of the cells identified by the S100AB antibody were also decorated by an antibody specific for S100B (Figure S1D), confirming the preferential reactivity of the S100AB serum for this latter molecule. S100A1+ cell of myelomonocytic origin, not stained by the S100B antiserum, were limited to mononuclear cells in the vessels, germinal center macrophages and terminally differentiated epithelial squamous cells (Figure S1D). The tonsil samples were serially stained with the antibodies listed in Table S1 and S2, the registered images were analyzed using the S100AB nuclear and cytoplasmic staining to produce a segmentation mask (henceforth named “S100-mask”). By computationally segmenting cells through this form of image analysis, the fluorescent intensities of all stains can be compared between selected cells. In order to assess the reproducibility of the analysis on manually selected areas and TMA cores, duplicate areas from each of the three tonsils and from three TMA cores were loaded in HistoCAT and analyzed by tSNE. Duplicates of the same sample superimposed, while subtle differences between tonsils for some of the phenoclusters appeared (not shown). Thus one single area of 3 mm^2^ or larger appear adequate to analyze tonsil tissue with a mask encompassing ~10% of its cell content.

Class II histocompatibility complex, HLA-DR, is considered a bona fide obligate marker for dendritic cells (DC), because of its requirement for antigen presentation; as a control for the representativeness of the S100-mask, a mask (henceforth named “DR-mask”) was prepared from an HLA-DR image from which CD20+ B cells were computationally subtracted. Differently from the S100-mask, segmentation of a membrane-based image is affected by the staining intensity and regularity of the membrane profile. Superimposition of the two masks on tissue images showed only a partial overlap.

In order to analyze identities and differences among the DC identified by the two masks, duplicate sets of images of the same field were loaded in HistoCAT, each with one mask, and tSNE and Phenograph analyses were run. Comparison of Phenograph clusters of cells segmented on the basis of S100AB or HLA-DR showed some superimposable clusters, some of partial overlap and some mutually exclusive clusters (Fig. 1). In general, S100-masks contained more cells than HLA-DR masks (Table 1 and Table S3).

**Figure 1.**
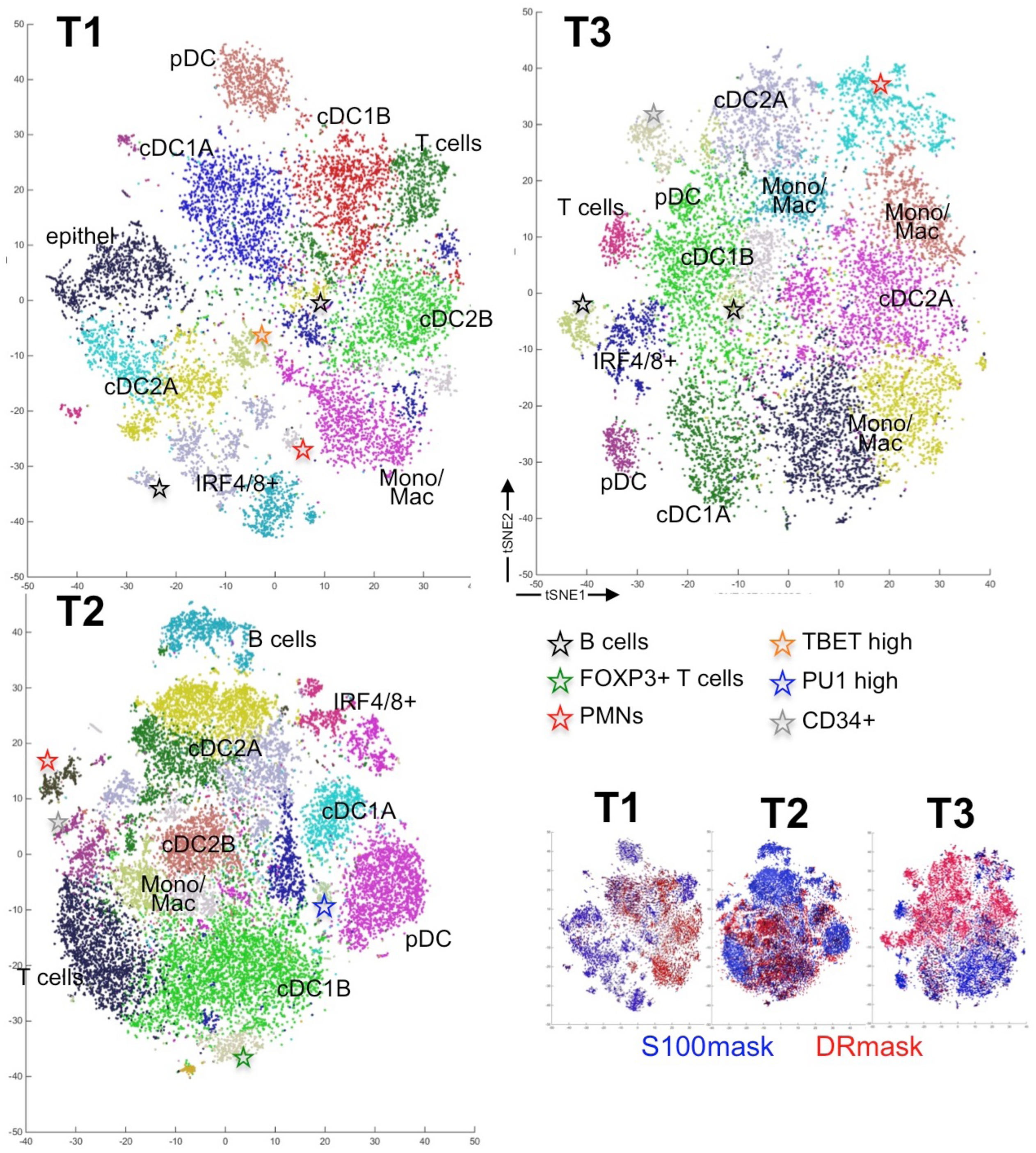
Annotated phenograph and tSNE plots for a selected area from each of three tonsils. Phenograph clusters are plotted on tSNE plots for each tonsil. The population name is written over the cluster/s in plain or as a colored asterisk. The legend for the asterisks is shown in the figure. The Phenograph settings were put at n=100. At the bottom right, the HLA-DR (red) and the S100AB (blue) masks are shown superimposed for each tSNE plot.

**TABLE 1.** Dendritic cell groups quantification in six tonsils.

| Phenogroups |  | on total DC* |  | on classic DC |  |
| --- | --- | --- | --- | --- | --- |
|  |  | mean | ±SD | mean | ±SD |
| **pDC** |  | **6.3%** | **±5.%** | - | - |
|  | pDC-A | 5.2% | ±5.% | - | - |
|  | pDC-B | 1.1% | ±2.% | - | - |
| **cDC1** |  | **20.6%** | **±3.%** | **28.0%** | **±9.%** |
|  | cDC1-A | 8.8% | ±4.% | 12.2% | ±7.% |
|  | cDC1-B | 11.8% | ±3.% | 15.8% | ±4.% |
| **cDC2** |  | **37.4%** | **±12.%** | **48.3%** | **±10.%** |
|  | cDC2-A | 21.8% | ±9.% | 28.5% | ±10.% |
|  | cDC2-B | 15.6% | ±10.% | 19.8% | ±13.% |
| **IRF4/8+ DC** |  | **9.5%** | **±5.%** | **12.5%** | **±6.%** |
|  | IRF4/8+-A | 6.7% | ±4.% | 8.5% | ±4.% |
|  | IRF4/8+-B | 2.9% | ±2.% | 4.0% | ±4.% |
| **MonoMac** |  | **23.8%** | **±11.%** | - | - |
| Total DC/segmented cells* |  | 80.5% | ±6.8% |  |  |
| Total classic DC/segmented cells |  |  |  | 61.6% | ±13.1% |
| Total segmented cells/DAPI |  | 13.2% | ±5.6% |  |  |

\* including MonoMac
Abbreviations: DC: dendritic cells; pDC: plasmacytoid DC; pDC-A: pDC expressing IRF8, TBET, MYC, pSTAT1, CD30 and very low levels of HLA-DR and IRF4; pDC-B: pDC with low levels of the markers expressed in pDC-A; cDC1: classic DC 1; cDC2: classic DC 2; IRF4/8+: DC coexpressing IRF4 and IRF8; MonoMac: monocytic/macrophagic cells; ROI: region of interests, individual cells digitally identified.

The cell type contained in each Phenograph group (phenocluster) was defined by four parameters: the location in the Phenograph (both absolute and relative to neighboring clusters), the distinguished phenotype, the composition in sub-groups by hierarchical clustering and the distribution in the tissue. Five main groups were identified in each of the six tonsil samples, four of them named similarly to previously identified DC subsets: 1) Classic DC1 (cDC1), 2) Classic DC2 (cDC2), 3) plasmacytoid DC (pDC), 4) “monocyte/macrophage” HLA-DR+ cells (MonoMac). A fifth group represents a novel putative DC subset co-expressing IRF4 and IRF8 (IRF4/8+ DC) observed here. The absolute and percentage numbers of these subsets are reported in Table 1 and Table S3.

80.5±6.8% of the cells sampled by both masks combined are hematopoietic cells of myeloid origin (cDC, pDC, MonoMac) and 61.6±13.1% are dendritic cells (pDC, cDC). cDC2 are the most represented DC in the tonsil (4.2±2.9% of all cells identified by DAPI), followed by MonoMac (2.2±0.9%), cDC1 (2.2±1.0%), IRF4/8+ DC (1.1±0.9%) and pDC (0.7±0.8%). Within the populations identified by the segmented cells of both masks, inclusive of the MonoMac cells, cDC2 is the most represented group (37±12%), followed by MonoMac (24±11%), cDC1 (21±3%), IRF4/8+ DC (10±5%) and pDC (6±5%).

### cDC1

Two major clusters observed to be nearby on Phenograph plots (Fig 1 and 2B), also occupy nearby microenvironments, i.e. T cell nodules between B cell follicles, closer to the stromal axis of the tonsil (Fig. 2C, 2D and Figure S4). One group, named cDC1-A, was identically represented by both masks, the other, cDC1-B, mostly from the DR-mask. Both groups shared expression of TIM3, but lacked CD1c, CLEC10A, CD123, IRF4 and myelomonocytic markers (CD14, CD16, CD68, CD163). cDC1-A, had prominent expression of markers associated with cDC1: IRF8, CD141, and high TIM3. In addition, cCD1-A expressed TF such as BCL6 (described in immature DC in the mouse (Zhang et al., 2014)), TBET, PU1, MYC, and lysozyme (LYZ), this latter often in a paranuclear Golgi location (Fig. 2E and Figure S2A, S3), and activation markers pSTAT1, CD30, PD-1. In tissue, cDC1-A was represented mostly by single, sparse cells (Fig. 2C and Figure S4).

**Figure 2.**
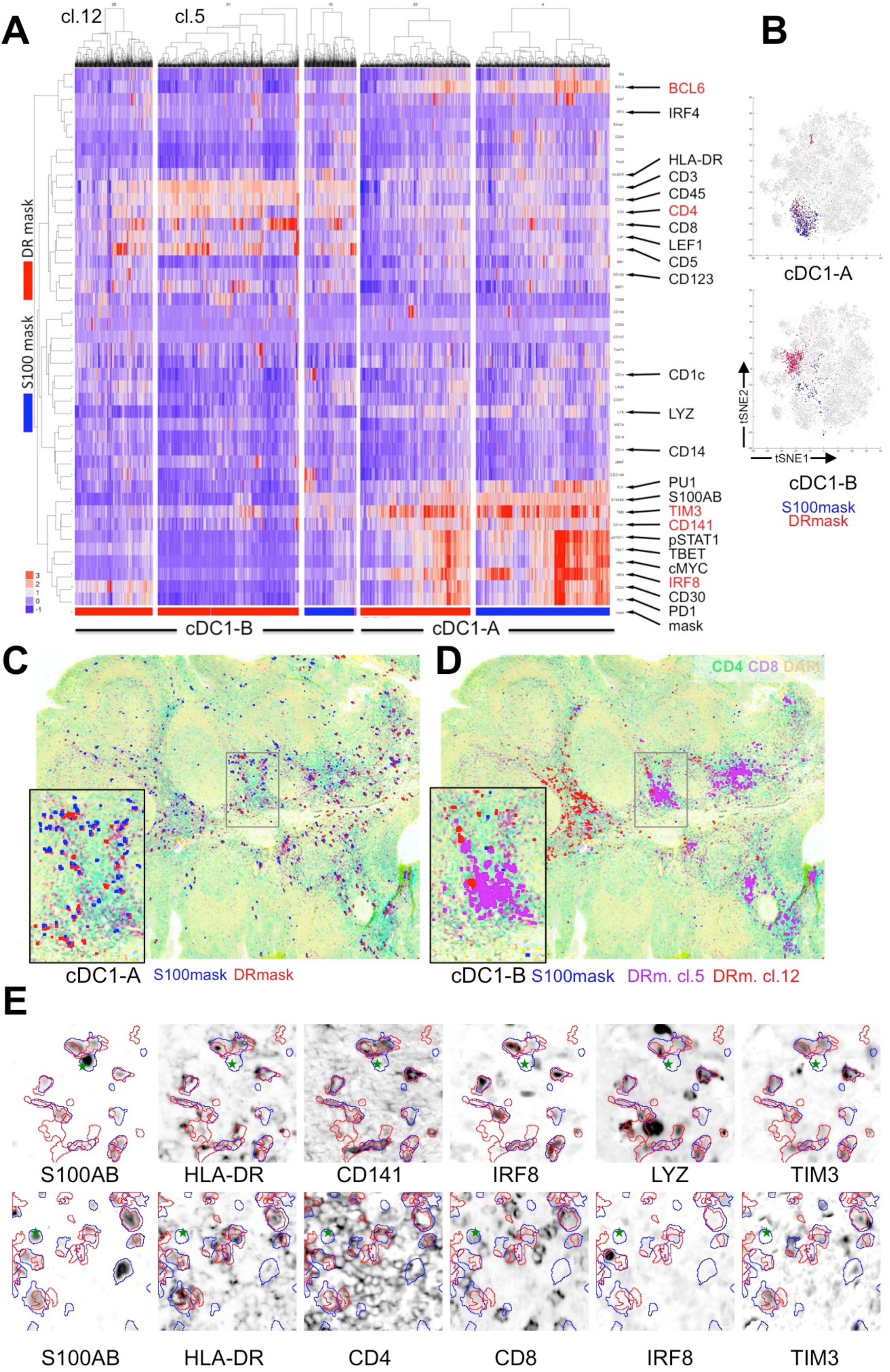
Classic Dendritic Cells 1 phenotype, clusters and tissue distribution. **A**: the five cDC1 Phenograph clusters from Tonsil 3 are shown, each one hierarchically clustered by markers (rows) and individual segmented cells (columns); the mask by which each phenocluster has been obtained is exemplified by the color band at the bottom: S100AB: blue, HLA-DR red. Two individual clusters in the cDC1-B group are indicated on top. Markers with arrows listed on the right are DC lineage and positive markers highlighted by the dendrograms. Distinctive and characterizing markers are in red. The two A and B subgroups are bracketed. Note in cluster 5 a cDC1-B population interacting with CD8+ T cells. **B**: The distribution of each phenocluster on the Tonsil 3 tSNE graph is highlighted with the dot color matching the mask color. **C**: The tissue distribution of the cDC1-A population is plotted against the tissue image of the tonsil from which the data were extracted. The tissue image is rendered by a subdued CD4, CD8 and DAPI staining. Each population is plotted with a color matching the mask from which the data are derived. An area highlighted by a rectangle in the main image is magnified in the inset. For image magnification see Fig S1. **D**: The tissue distribution of cDC1-B population is plotted as in **C**. The two clusters identified in **A** are plotted in contrasting colors. An area highlighted by a rectangle in the main image is magnified in the inset. For image magnification see Fig S1. **E**: TOP row: five images of selected stains, representative of the cDC1-A population, are shown as inverted fluorescent images; the color-coded outline of the S100AB and DR masks are superimposed. The stain name is written below each image. Images size 89×89 μm. BOTTOM row: five images of selected stains, representative of the cDC1-B population, are shown as above. Images size 89×89 μm.

About a third of the cDC1-A cluster showed greatly reduced intensity of expression for TBET, MYC, BCL6, pSTAT1, PU1 and activation markers, less so for IRF8, TIM3 and CD141 (Fig. 2A and Figure S2A), a phenotype found in all cDC1-B cells. These latter cells featured in addition an evident T cell signature, composed predominantly of CD4+ T cells and a minor, heterogeneously represented CD8+ subset (Fig. 2A and Figure S2A). By single cell inspection, the T cell signature was due to a close interaction of cDC1-B with T cells and pixel overlap, detected by both masks on cells closer than 1 μm (2 pixels), and not an autonomous DC phenotype (Fig. 2E and Supplementary Fig.3). The cCD1-B cells had prominent aggregation in the same T cell areas shared with cDC1-A cells (Fig. 2C, D and Figure S4), although not occupying the same identical microscopic location. T-cell interacting cDC1-B displayed subtle phenotypic differences, picked up by the Phenograph algorithm (Fig. 2A; clusters 5 and 12) and spatially resolved in the tissue as distinct interfollicular zones separated one B cell follicle apart (Fig. 2D). The downregulation of the TF expression was a DC-autonomous phenotype, not the dilution of the signal by neighboring cells (Fig. 2E and Figure S3).

### cDC2

Multiple phenoclusters in both masks display clearly distinct Phenograph regions (Fig. 1 and 2C) and show evidence of cDC2 phenotype (HLA-DR, CD1c, CLEC10A, LYZ) in the absence of cDC1, myelomonocytic and plasmacytoid DC markers (Fig. 3A, D and Figure S2B, 5). Differently from the cDC1, the clusters in this group were quite variegated, in both masks. One group, identified by the S100-mask and named cDC2-A was characterized by an intra- and sub-epithelial location toward the surface of the tonsil (Fig. 3B and Figure S4), a B cell signature caused by a close association with intraepithelial B cells but not follicle mantle or germinal center B cells (Fig. 3A, D and Figure S5), interferon associated signaling (MX1, pSTAT1, TBET), absence of IRF4, IRF8, LYZ and partial CD16 expression. The cDC2-B group had an interfollicular distribution, toward the axis of the tonsil (Fig. 3B and Figure S4) and displayed CD1c and CLEC10A, sometimes in an uncoordinated fashion, pSTAT1, low and variegated expression of MYC, CD1a, CD207, CD16, CD14, VISTA, and LYZ (Fig. 3A and Figure S5). T cell-related, macrophage, pDC markers and IRF4 were absent.

**Figure 3.**
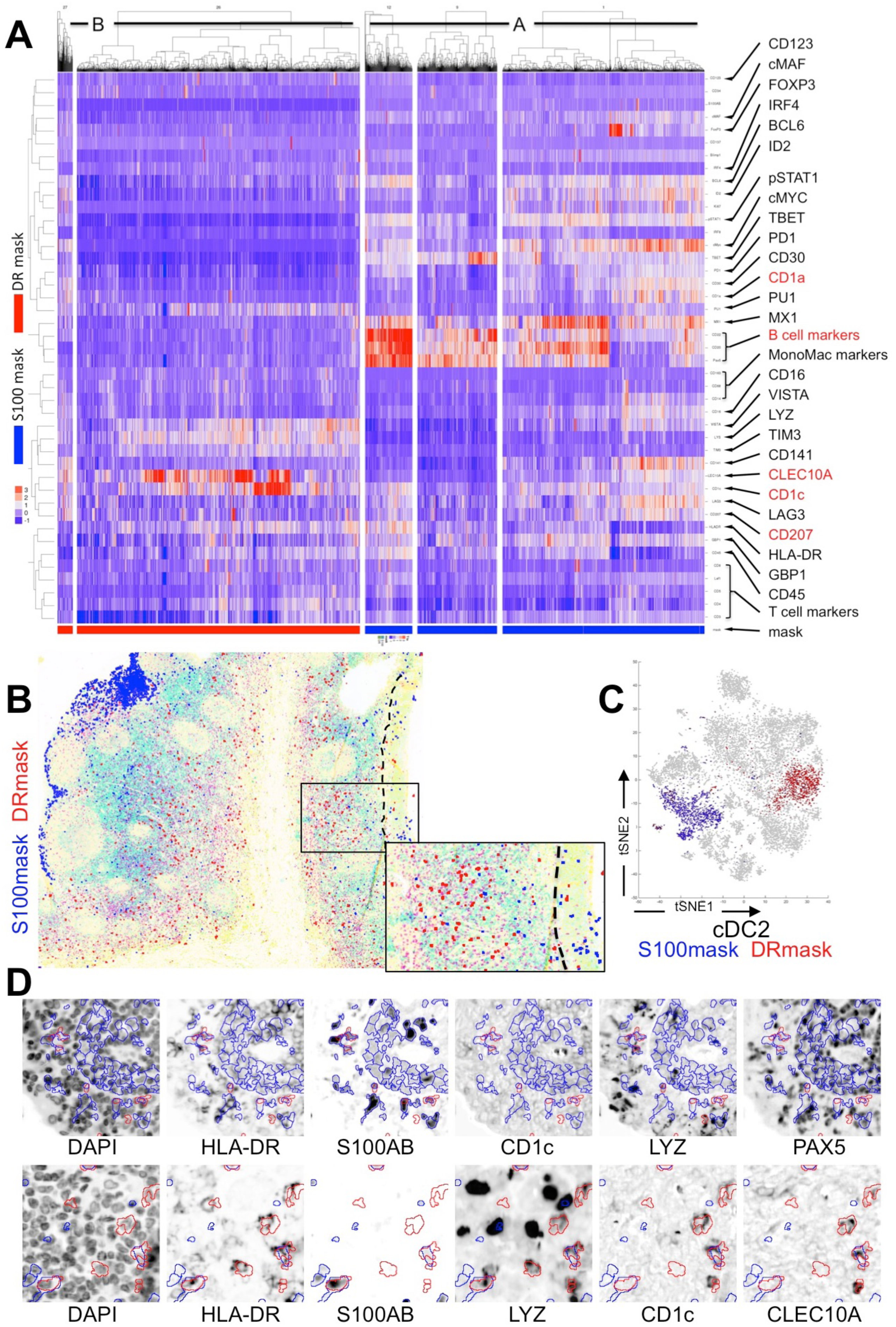
Classic Dendritic Cells 2 phenotype, clusters and tissue distribution. **A**: the five cDC2 Phenograph clusters from Tonsil 1 are shown, as per Fig.2A. The A and B subgroups are bracketed. **B**: The tissue distribution of the cDC1-A population is plotted against the tissue image of the tonsil from which the data were extracted. The tissue image is rendered by a subdued CD4, CD8 and DAPI staining. Each population is plotted with a color matching the mask from which the data are derived. The dashed line highlights the squamous epithelial basal layer. An area highlighted by a rectangle in the main image is magnified in the inset. For image magnification see Fig S1. **C**: The distribution of each phenocluster on the Tonsil 1 tSNE graph is highlighted with the dot color matching the mask color. **D**: TOP row: five images of selected stains, representative of the cDC2-A population, are shown as inverted fluorescent images; the color-coded outline of the S100AB and DR masks are superimposed. The stain name is written below each image. Images size 118×118 μm. BOTTOM row: five images of selected stains, representative of the cDC2-B population, are shown as above. Images size 89×89 μm. See also Fig. 1, S2B, S4 and S5.

### IRF4/IRF8+ DC

In each of the six tonsil samples, a small phenocluster (Fig. 1 and 4B) was identifiable by its co-expression of IRF4 and IRF8, albeit in part of it (Fig. 4A, D and Supplementary Fig.2D, 3). The cells belonging to this group, named IRF4/8+ DC, reside between the follicles, spread upward toward the epithelium and downward in the stromal axis. Besides the two TF named above, this group coordinately express ID2, PRDM1, TBET, MYC, pSTAT1 and lacks T-, B-, cDC1, cDC2, pDC and myelomonocytic lineage markers (Fig. 4A, D and S2D, S3). Ki-67 separate the IRF4/8+ DC into two subpopulations, IRF4/8+ DC-A and −B. The Ki-67 negative subpopulation, IRF4/8+ DC-B, often expresses VISTA and positions itself in the surface subepithelial layers (Fig. 4C and Figure S4). In tonsil T1, the IRF4/8+ DC-A cluster contained a group of interfollicular proliferating B cells coexpressing IRF4, IRF8 and PRDM1, clearly identifiable by intra-phenocluster hierarchical clustering (Figure S2D). No close interaction with T or B cells or macrophages was noted however.

**Figure 4.**
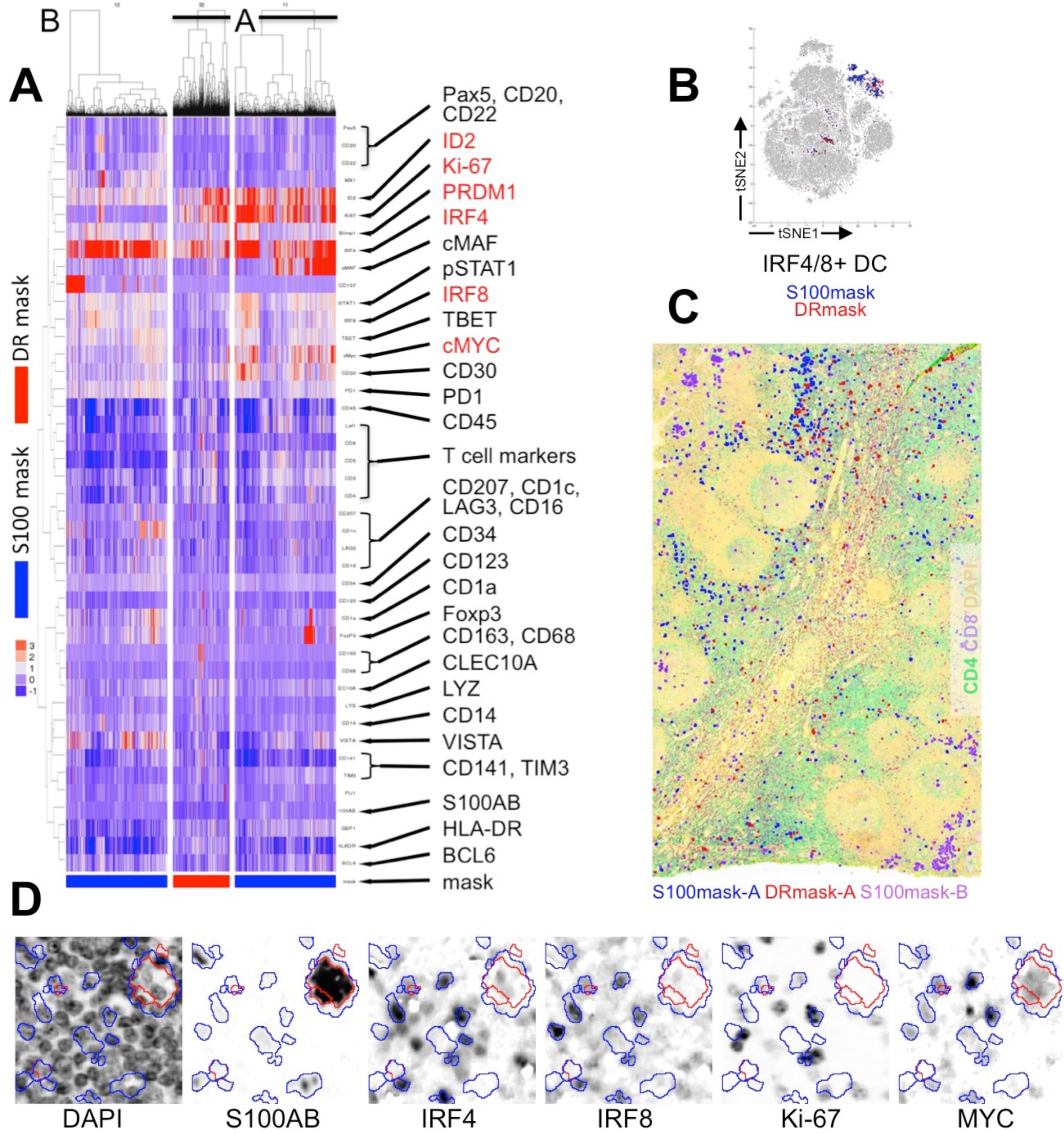
IRF4 and IRF8 coexpressing Dendritic Cells phenotype, clusters and tissue distribution. **A**: the three IRF4/8+ DC Phenograph clusters from Tonsil 2 are shown, as per Fig.2A. The A and B subgroups are labeled. **B**: The distribution of each phenocluster on the Tonsil 2 tSNE graph is highlighted with the dot color matching the mask color. **C**: The tissue distribution of the IRF4/8+ DC population is plotted against the tissue image of the tonsil from which the data were extracted. The tissue image is rendered by a subdued CD4, CD8 and DAPI staining. Each population is plotted with a color matching the mask from which the data are derived. **D**: five images of selected stains, representative of the IRF4/8+ DC population, are shown as inverted fluorescent images; the color-coded outline of the S100AB and DR masks are superimposed. The stain name is written below each image. Images size 70×70 μm. See also Fig. 1, S2D, S3 and S4.

### pDC

One group of cells was the most recognizable in each Phenograph because of its compactness and separation from the bulk of clusters (Fig. 1 and 5B): it was composed of CD123+ IRF8+ CD68+ cells and we assigned it to the plasmacytoid DC lineage (Fig. 5A, D and Figure S2E, 5). The proportion of pDC identified by the S100-mask vs the DR-mask was very variable and sample-dependent. In both masks, pDC occupied the lymphoid tissue between the area containing the follicles and the stromal axis (Fig. 5D and Figure S4). The DR-mask comprises pDC aggregates. A major phenotypic difference between the pDC identified by the S100AB and the DR-masks was that the former displayed abundant signal for IRF8, TBET, MYC, pSTAT1, CD30 and very low levels of HLA-DR and IRF4 (Fig. 5A, D and Figure S2E and 5). CD123 and CD68 remained invariant across the two masks. An additional population of CD123+ lymphoid cells, lacking HLA-DR, S100AB, IRF8, thus not comprised by the masks used was identified on the images (Fig. 5D). To independently confirm the phenotype of the pDC, these were analyzed in the same areas with a mask comprising all CD123+ cells, including positive high endothelial venules (HEV). tSNE analysis of the CD123+ cells showed very little overlap of the IRF8+ and the HLA-DR+ populations (Fig. 5E). Bivariate plots of IRF8, pSTAT1 and CD68 (as a control) against HLA-DR showed inverse relationship between IRF8, pSTAT1 and HLA-DR (Fig. 5F), confirming the observation made with the S100 and DR masks. The CD123 mask, once the HEV were removed by CD34 image subtraction, yielded 75.5%±50.0% more pDC than the combined S100 and DR masks, placing this subset fourth by abundance among DCs.

**Figure 5.**
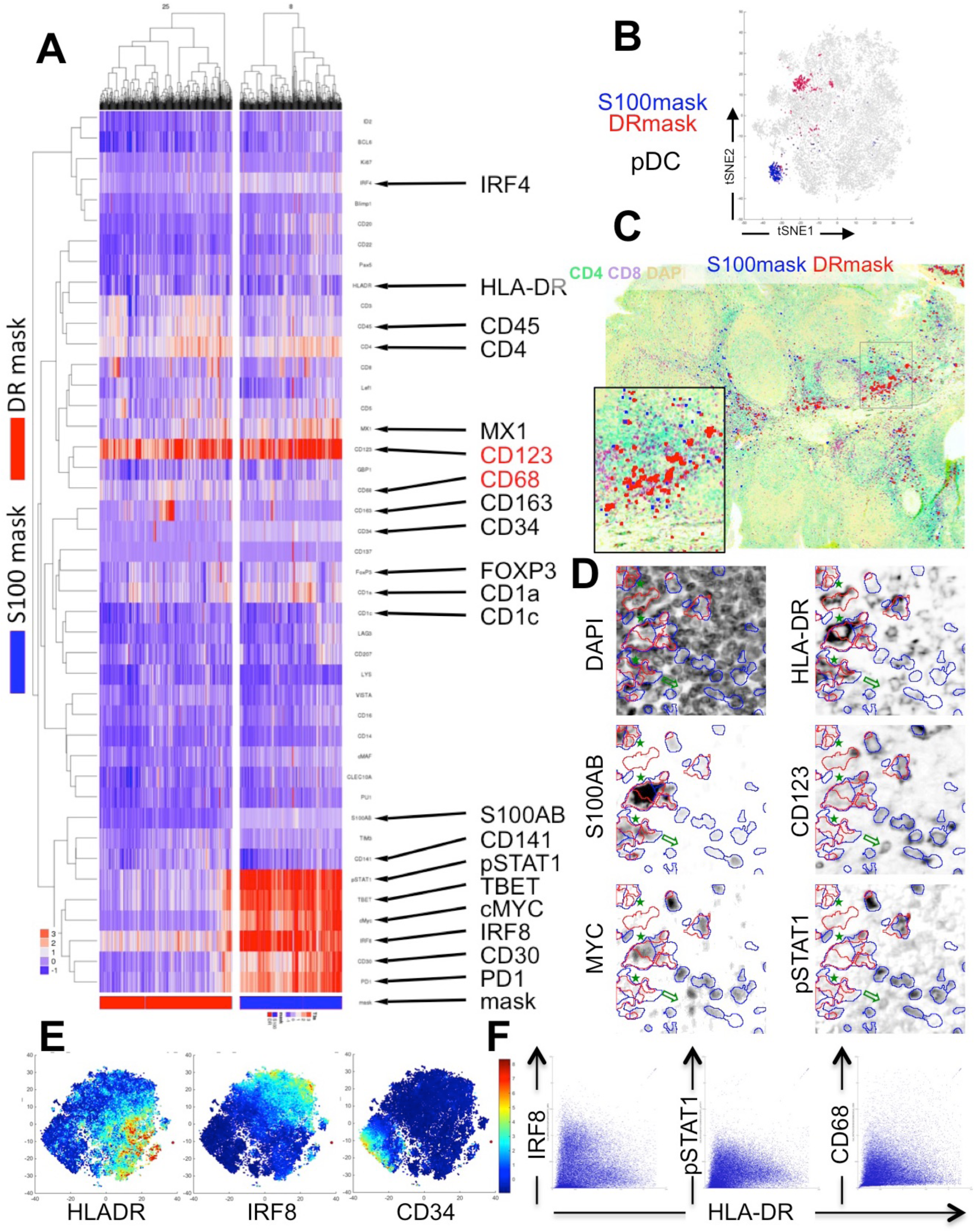
Plasmacytoid Dendritic Cells phenotype, clusters and tissue distribution. **A**: the two pDC Phenograph clusters from Tonsil 3 are shown, as per Fig.2A. **B**: The distribution of each phenocluster on the Tonsil 3 tSNE graph is highlighted with the dot color matching the mask color. **C**: The tissue distribution of the pDC population is plotted against the tissue image of the tonsil from which the data were extracted. The tissue image is rendered by a subdued CD4, CD8 and DAPI staining. Each population is plotted with a color matching the mask from which the data are derived. An area highlighted by a rectangle in the main image is magnified in the inset. For image magnification see Fig S1. **D**: five images of selected stains, representative of the pDC population, are shown as inverted fluorescent images; the color-coded outline of the S100AB and DR masks are superimposed. The stain name is written below each image. Three non-pDC cells can be seen on the left (stars). A CD123+ cells outside the masks is highlighted by a green arrow. Images size 84×84 μm. **E**: tSNE plots of cells segmented with a CD123+ mask, colored for HLA-DR, IRF8 and CD34. Note the minimal overlap of HLA-DR and IRF8 and the separation of the CD34+ endothelial cells. **F**: bivariate plot of cells segmented with a CD123+ mask, showing inverse relationship between IRF8, pSTAT1 and HLA-DR. CD68 is shown as a control. See also Fig. 1, S2E, S4 and S5.

### DR+ cells

**Monocytic/macrophagic HLA**

A large and heterogeneous collection of phenoclusters, largely represented by the DR-mask is marked by the variegated and uncoordinated expression of myelomonocytic markers (CD14, CD16, CD68, CD163, lysozyme) but none of the prototype cDC1, cDC2 or pDC markers (Fig. 6A, D and Figure S2C). PU1 and cMAF, known to be expressed in myelomonocytic cells (Bakri et al., 2005; Barros et al., 2013), were detected. None of the TF expressed in DC and linked to IFN γ signaling was consistently present except GBP1. Guanylate-binding protein 1 (GBP1) is a protein highly induced by IFNγ (Lubeseder-Martellato, 2003), but we found it constitutively expressed in myeloid cells.

**Figure 6.**
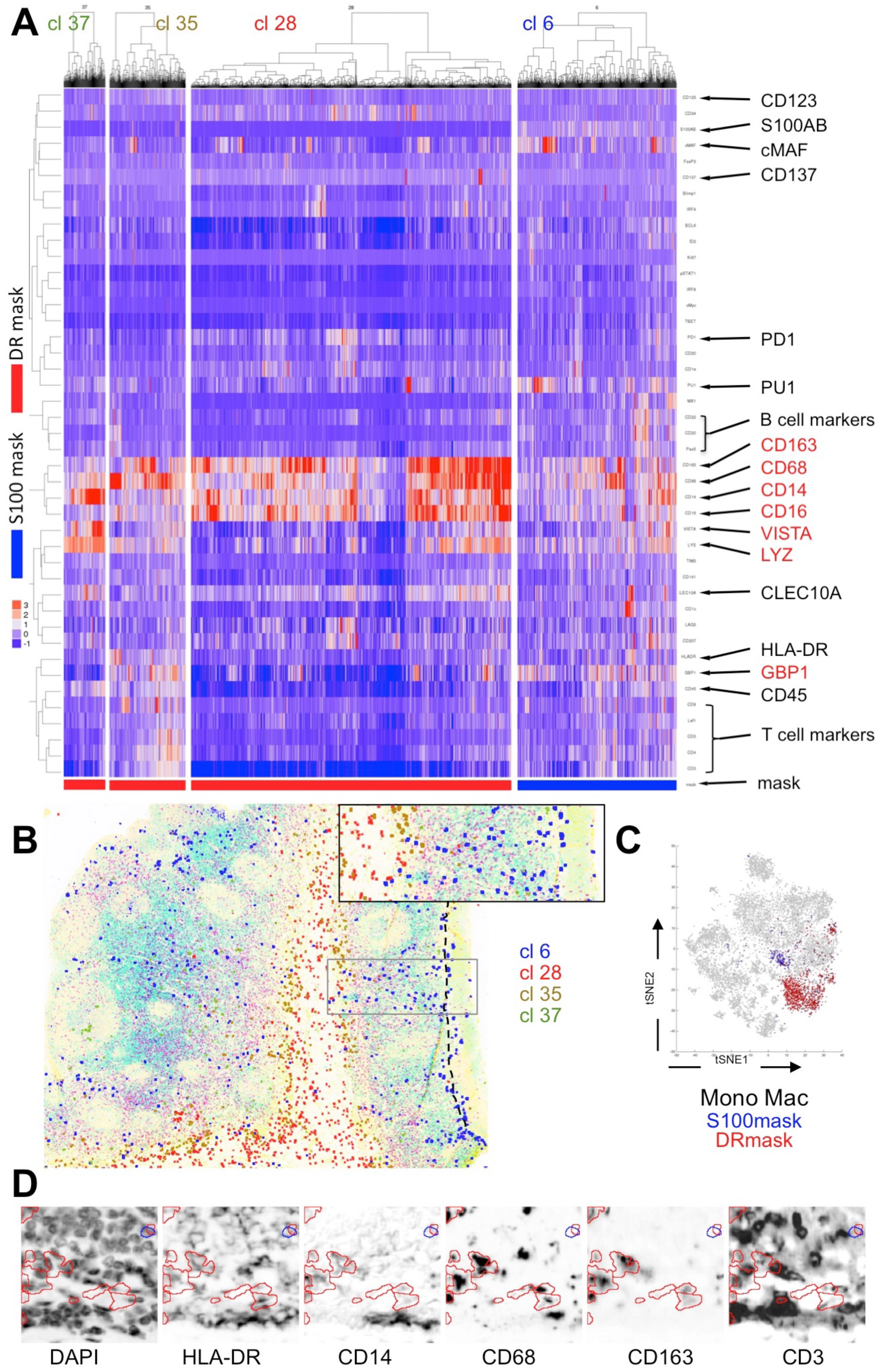
Monocyte Macrophage phenotype, clusters and tissue distribution. **A**: the four MonoMac Phenograph clusters from Tonsil 1 are shown, as per Fig.2A. **B**: The tissue distribution of the MonoMac population is plotted against the tissue image of the tonsil from which the data were extracted. The tissue image is rendered by a subdued CD4, CD8 and DAPI staining. Each population is plotted with a color matching the phenocluster from which the data are derived. The dashed line highlights the squamous epithelial basal layer. An area highlighted by a rectangle in the main image is magnified in the inset. For image magnification see Fig S1. **C**: The distribution of each phenocluster on the Tonsil 1 tSNE graph is highlighted with the dot color matching the mask color. **D**: five images of selected stains, representative of the MonoMac population, are shown as inverted fluorescent images; the color-coded outline of the S100AB and DR masks are superimposed. The stain name is written below each image. Images size 75×75 μm. See also Fig. 1, S2C and S4.

Subsets of MonoMac cells in the S100-mask, in numbers in excess of the S100A1+ cells (Figure S1D) may represent activated histiocytes, which have been shown to express S100B upon IFN and IL4 signaling (Riuzzi et al., 2017). The tissue localization of the MonoMac phenocluster was along the stromal axis, on the epithelial surface or in an uncharacterizing sparse location (Fig. 6B and Figure S4). However, the numerosity varied among samples and there were hardly a common phenotypic group shared by two samples. There seemed to be no interaction with T, B lymphocytes or the previously described DC subsets.

A separate group of DC, named inflammatory DC (Segura and Amigorena, 2013) has been described, resembling in vitro differentiated DC from peripheral monocytes. The signature of inflammatory DC is close to cDC2 and monocytes (Segura and Amigorena, 2013). We could not find a similar cell in our samples, the closest being CD16+ cDC2-A, which however do lack other myelomonocytic markers.

### T cells engaging cDC1

In order to understand the functional implications of the T cell-DC contact, a mask was created with the T-cell specific nuclear TF LEF1. tSNE and Phenograph analysis revealed T cell phenogroups with largely overlapping distribution in the tissue section (Fig. 7E and not shown). Of the groups representing several individual or relation-dependent phenotypes of T cells we selected two: one containing T cells interacting with cDC1-A (Fig. 7A and C) and one with pDC. (Fig. 7B and D). In both groups the phenotype of the T cells was exclusively dictated by the encroaching DC type, mirroring what has been described above for cDC1 and cDC2. Early activation (CD137, IRF4) and proliferation markers were absent.

**Figure 7.**
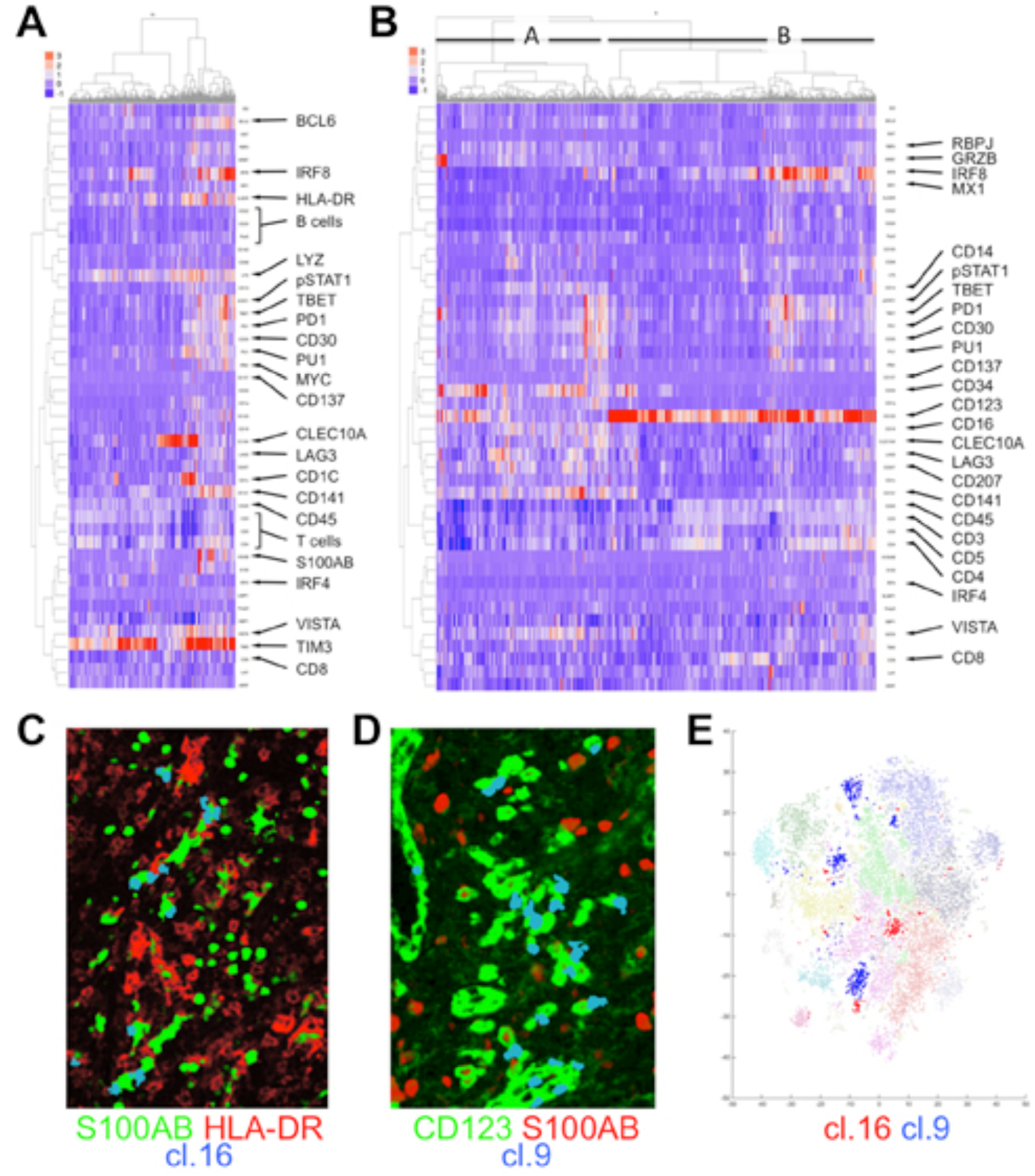
DC-interacting T cell populations phenotype and tissue distribution. **A:** heatmap of cluster 16, interacting with cDC1-A is shown, as per Fig.2A. **B:** cluster 9 interacting with pDC (B) and trafficking across the CD34+ CD141± high endothelial venules (HEV; A) is shown, as per Fig.2A. **C:** plotting of the phenocluster 16 as single cells shows T cells (blue) touching S100+ (green) cDC1-A (verified by plotting both T and DC clusters), in an area containing HLA-DR+ cDC1-B with elaborate dendrites (red). **D:** plotting of the phenocluster 9 as single cells shows a pDC-rich area where T cells (blue) touch CD123+ (green) pDC, both S100+ and S100-, and cross CD123+ HEV (bottom). **E:** the phenoclusters 16 (blue) and 9 (red) are highlighted over a Phenograp map of all LEF1 T-cell clusters.

Although other T cells phenogroups specifically interacted with DC, e.g. a FOXP3+ T cell group interacting with cDC1-B or a cDC2 interacting group (not shown), the tissue distribution was broader, thus not informative of the specific interaction.

### Other Identified Cell Populations

In addition to the established DC and monocyte/macrophages groups discussed above, our two mask approaches also identified S100+ HLA-DR- CD45+ CD8+ and CD4+ T cells (Figure S4 and not shown), a subset of activated HLA-DR+ T cells, partially expressing FOXP3 (Figure S6A) and TBET (Figure S6B), PU1 and S100AB overexpressing cells (Figure S6C) of uncertain lineage. In a single tonsil, S100AB+ CD14+ follicular dendritic cells coopted Germinal Center B cells (BCL6+, Ki-67+, CD14-) by contiguity in the mask (Figure S6A). Furthermore, we identified phenoclusters of CD45- HLA-DR-S100A1+ epithelial cells (Figure S6D) and endothelial cells (Figure S6E, F, G).

## DISCUSSION

Dendritic cell subsets have been previously identified in cell suspension by traditional flow cytometry, single cell RNA sequencing or ion beam flow cytometry such as CYTOF (Alcántara-Hernández et al., 2017; Guilliams et al., 2016). We present here an in situ high-parameter analysis via a robust and versatile technique. The in situ approach has allowed us to reproduce and refine most published DC subsets, but has added a detailed topographic information on the distribution of DC subsets in the tonsil microenvironment, insights into their preferred cellular partners by their close engagement and changes in the TF repertoire after cell-cell communication.

We examined 6 tonsils with two approaches to image analysis by object segmentation, using either a HLA-DR and S100AB masks, which overlap by an average of 53% and together sampled 13% of tonsil tissue cells. Differently from published experiments that examined cell suspensions, we excluded from the analysis only HLA-DR+ CD20+ B cells by image subtraction, allowing the sampling of hematopoietic cells of dendritic, myelomonocytic and T lineage and of non hematopoietic lineage (vascular, epithelial, stromal). We found that pDC do express S100B, validating it as a “pan-dendritic” marker.

Our data on intact tissue show that cDC2 are the most represented DC in the tonsil, followed by Monocytes/Macrophages (MonoMac), cDC1, IRF4/8+ DC and pDC. cDC1 is the subset with the least variation in numbers across the six samples, consistent with a tissue-patrolling function that may be less influenced by the activation status of the tonsil tissue in which resides. Comparable quantitative data in literature are scarce: we compared data from tissue disaggregation reported by six groups (Alcántara-Hernández et al., 2017; De Monte et al., 2016; Granot et al., 2017; Heidkamp et al., 2016; Jongbloed et al., 2010; Segura et al., 2012) and found that cDC and pDC frequencies for tonsil tissue or mucosa-associated lymphoid tissue were underreported in each instance, occasionally severalfold less. cDC1 data ranged from one fourth to fortyfold lower, cDC2 from half to eightfold less. pDC numbers instead ranged from twice to ten times higher the figures we have found. Disaggregating a piece of lymphoid tissue to obtain a single cell suspension poses a risk of selectively losing in the process intrinsically fragile cells such as plasma cells or cells with irregular, dendritic extensions, such as DCs. Furthermore, DCs which are often enveloping other cells with their dendrites, because of physiologic interaction, may be lost by doublet exclusion during flow cytometry analysis. Both mechanisms may occur during tonsil disaggregation and may selectively affect DC subpopulations that are prone to interact with other lymphoid cells such cDC1-B, cDC2-A and pDCs captured by the DR-mask and found to aggregate. pDC are closer in shape to lymphoid cells and may be relatively enriched by the process. The inverse cDC/pDC ratio and the broader variations across the mean reported by the papers cited above suggests a selective cDC loss during tissue disaggregation, particularly the cDC1 group.

All DC and MonoMac populations identified express HLA-DR to various extents and DR is among the markers with the broader differential between positive and negative cells in tissue (Fig S1E). However, besides cDC1 and the IRF4/8+ DC population, where the S100-mask and the DR-mask largely coincide, the two masks seldom overlap. Furthermore, they seem to highlight a phenotypic difference with functional, maturation or topographic implications. We found the S100-mask to be very effective at capturing tissue cells with irregular dendritic contours. By inspecting high-magnification images, cells bearing morphological features or stereotypical surface markers for DC were consistently captured by either this or the DR-mask. Despite the fact that we report the highest number of DC published so far, we cannot exclude that a subset of DC may have gone undetected by our approach; in fact we did observed CD123+ cells not captured by either mask.

There is little information on the role of S100B in dendritic and myeloid cells. S100B conformation is sensitive to Ca^2+^ ions and targets numerous proteins in a broad range of cell types and metabolic pathways, including those that affect heart, skeletal and cartilage tissues. (reviewed in (Donato, 1999)). For example, secreted S100B affects skeletal muscle repair and favors M1 to M2 macrophage transition and migration (Riuzzi et al., 2017). In this setting, macrophages acquire de-novo S100B via an NF-kB-dependent IFNγ stimulation and -independent IL-4 stimulus (Riuzzi et al., 2017). Notably, the S100+ DC phenoclusters show the expression of several TFs, including TBET, phospho STAT1 and MX1, all associated with IFN signaling (Hu and Ivashkiv, 2009; Lugo-Villarino et al., 2003; Simon et al., 1991). TBET has been shown to be central for an autocrine, STAT1-dependent IFNγ induction loop in T cells (Lighvani et al., 2001). TBET in DCs is required to boost Th1 IFN-dependent T cell immunity (Lugo-Villarino et al., 2003). Although myeloid cells may be induced to synthesize IFNγ, it is unclear if DCs have the ability. In summary, S100B+ DC, independently of the inclusion in the DR-mask, seems to show evidence of IFN signaling and activation.

MYC, which is selectively expressed in the cDC1-A, cDC2-A, pDC-A and IRF4/8+ subsets (Heidkamp et al., 2016), may confer metabolic adaptation (Kress et al., 2015; Stine et al., 2015) and plasticity in lineage differentiation (Wilson et al., 2004a). Within the hematopoietic system, MYC upregulation is required for the transition of stem cells from the quiescent status in the niche into the transit amplifying compartment (Ehninger et al., 2014; Murphy et al., 2005) while maintaining an undifferentiated state. Many different mitogenic stimuli cause a rapid MYC upregulation via changes in protein stability (Ehninger et al., 2014; Eisenman, 2001). IFN signaling is a likely candidate for MYC upregulation in DC, given the coexpression of IFN-related molecules in the same cells. It is intriguing to observe the selective MYC expression in cells expressing S100B; reminiscent of the ability of MYC to prevent a terminally differentiated phenotype (Lin et al., 2000; Shachaf et al., 2004) is the capability of S100B and S100A1 to produced a poised intermediate state of differentiation upon SOX TF signaling, albeit in a mesenchymal setting (Saito et al., 2007).

Immature DC have reduced total and surface amounts of HLA-DR (Cella et al., 1997; Granot et al., 2017). DC identified by a robust, even surface display of Class II antigens, thus may represent a more mature state. The data presented here suggests that for DC groups sharing an unique tissue niche such as cDC1 and pDC, there is a transition from a predominantly S100B+ status to an HLA-DR^high^ status. Along this transition, DC go from single dispersed cells (e.g. cDC1-A, pDC-A) to cells found in aggregates, such as cDC1-B and pDC-B. For cDC1, the close interaction with T cells seems to be the driving stimulus for a substantial phenotypic change, as shown. Here we report for the first time a cDC-T cell interaction with a coordinate and distinct transcription factor profile in vivo in human lymphoid tissue. This was not an unexpected finding given that previous experimental data published in mouse and ex vivo in humans, these latter based on cell suspensions (Ingulli et al., 1997; Itano et al., 2003; Krishnaswamy et al., 2017; Mempel et al., 2004; Miller et al., 2004). DCs which specifically interact with antigen-reactive T cells have been reported to reduce motion and increase the contact time with cells of cognate specificity and cluster in situ (Gerner et al., 2017; Mempel et al., 2004); this may translate into the aggregated form we have documented in tissue. We can add to this the reduction of markers of endocytic trafficking (LYZ) and of cognate inhibitory signals (TIM3) and the loss of IFN signaling related proteins (pSTAT1, TBET). Differently from an analysis of the transcriptional consequences of DC-T interactions in a murine model (Pasqual et al., 2018), we did not observed a reduction in MX1 protein in DC after a contact with T cells, possibly because we monitored post-translational changes. We were unable to discern the stimulus driving pDC transition from an −A to a −B state, despite the fact that pDC have the most dramatic fluctuations between the two subsets in all tonsils.

While we could document for the first time a variegation of the phenotype of DC, particularly for TF, we searched for a phenotypic evidence of the T cell-DC interaction in two T-cell subsets in teracting with cDC1-A and pDC, but found none. The relative limitation of the immunopanel employed may be one reason. The other, the contact time for T-DC interaction is limited to <72 hrs, after which T cells, extremely motile, travel beyond the few microns distance from the interacting partners, escaping our analysis of fixed tissue and coming close to partners with whom they have transient, non physiologic interactions (Pasqual et al., 2018), blurring the picture. Thus, the effect of the interaction resulting in a change in protein repertoire occurs when the T cell is no longer available to our analysis based on spatial distribution.

cDC2, the largest DC population in tonsils, is split into two subgroups by the tissue location and by interaction with B cells. The expression in cDC2-A of CD16, LYZ, VISTA, BCL6, and, limited to S100B+ subset, of TBET, MYC and pSTAT1 suggests that DC interacting with B cells differ from the ones that don’t. The expression of prototypic markers for cDC2 (CLEC10A and cDC1c) is asynchronously distributed across the two subgroups. We could not confirm a substantial presence of IRF4 in cDC2, differently from others who detected this TF in humans (Granot et al., 2017) and in both human and mouse (Guilliams et al., 2016). Memory B cells (Liu et al., 1995) lodge within the tonsil reticulated epithelium, which were not directly stained in these experiments. We cannot exclude that cDC2-A are in fact interacting with the epithelium rather than with B cells, however the presence of a pixel overlap with adjacent B cells is consistent with an interaction with B cells. DC interaction with B cells has been previously described in tissues (Takahashi et al., 2006) and in vitro experiments have shown a T-independent B cell activation (Balazs et al., 2002; Chappell et al., 2012; Qi et al., 2006). Consistent with this, murine CD8^neg^ DCs, the equivalent of cDC2, have the ability to induce B cell activation (Balazs et al., 2002).

The unsupervised analysis revealed in every sample a consistent group of IRF4/8+ proliferating cells that were not lymphocytes. This subset co-expressed TF associated with DC differentiation (ID2, MYC), IFN signaling (TBET, pSTAT1) and IFN autoregulatory loops (PRDM1) (Cimmino et al., 2008). None of the DC, pDC and myelomonocytic markers previously known or discovered during this investigation were expressed by the proliferating IRF4/8+ cells. However, ID2 and PRDM1, two TF known to be essential for DC differentiation and maturation (Hacker et al., 2003; Smith et al., 2011; Watchmaker et al., 2014), were uniquely expressed in IRF4/8+ DC but not in other subsets. Because the set of TF expressed by these cells have been described as crucial for DC differentiation (Hacker et al., 2003; Lee et al., 2017) and expressed in pre-DC (See et al., 2017), we explored their relationship to the dendritic lineage. The tissue localization of this subset is distant from the areas occupied by cDC1, cDC2, pDC and MonoMac. This group of cells may therefore represent precursors seeding the tonsil and proliferating before they differentiate. Low levels of CD34 we found reinforce this hypothesis. Notably, none of the other phenoclusters described has any significant proliferating fraction. A subset of this IRF4/8+ DC subset is not proliferative: these are often VISTA, CD1a positive and are allocated in part in the subepithelial regions, where cDC2-A reside.

AXL+ DC with a phenotype intermediate between pDC and cDC2, expressing TCF4, CD5 and CD2 have been described (Alcántara-Hernández et al., 2017; See et al., 2017; Villani et al., 2017). The nature of these cells is debated (Alcántara-Hernández et al., 2017). RNAseq and CYTOF experiments (See et al., 2017; Villani et al., 2017) suggests that these cells are pDC precursors; gene expression analysis indicate the presence of CD22 and CLEC10A (Heidkamp et al., 2016; See et al., 2017), which we tested. In tonsil suspensions, Axl+ DC amount to 3% of total Lin-DR+ cells, being cDC1 4%, cDC2 15% and pDC 78% in the same experiments (Alcántara-Hernández et al., 2017). We could not identify any subset corresponding to this population: CD5 and CD22 signal belong invariably to T and B cells encroaching the mask and carried the full antigenic repertoire of the cell of origin (i.e. CD3, LEF1, PAX5, CD20 etc.). This is surprising in view of the substantial proportion of these cells in tonsils reported (Alcántara-Hernández et al., 2017), however a relative enrichment of very very rare blood-borne cells because of a dramatic loss of other more abundant cells (e.g. cDC1) during disaggregation may explain the discrepancy.

Our in-situ approach benefits from the identification of other cell types at a micron’s distance. DCs of the cDC1 subset can differ in cell-autonomous expression patterns if they are in contact with a T cell. Of note, we could not observe DC-macrophage interaction to any extent, suggesting that the transfer of antigen from a phagocyte to a DC does not occurs, or not in the tonsil but elsewhere, or via such a limited and transient contact to escape our in-situ analysis. The study presented here has cataloged all DC subsets in situ in whole tonsillar tissue and further identified their preferred lymphocyte interactions in the context of tissue architecture.

Preliminary data on lymph nodes hosts of immune activation suggests that this high-dimensional approach can unveil additional tissue-specific heterogeneity in DC, depending of the lymphoid organ (nodal vs extranodal vs mucosal) and the prevalent type of immune activation (paracortical vs follicular hyperplasia), an area we are currently investigating with an appropriate selection of donors/sites.

The tissue map of DC, defined by phenotype, microenvironmental location and cell interaction, provides a discrete, well defined repertoire of cell types, on which experimental approaches may build more dynamic observation. Using this powerful approach, other meaningful immune cell interactions could be likewise explored in future studies of other tissue types in health and disease.

## ACKNOWLEDGEMENTS

We wish to thank Dr. Franco Ferrario for continuous support, Emanuele Martella (Nikon, Italia) and Giulio Simonutti (Hamamatsu Italia) for expert advice. The Hamamatsu S60 digital scanner was obtained as part of a clinical research project BEL114054 (HGS1006-C1121) of the University of Milano-Bicocca and GlaxoSmithKline, on which project Maddalena Maria Bolognesi is also supported.

This work has been supported by Departmental University of Milano-Bicocca funds. RT and MMB are employed by the Department of Medicine and Surgery of the University of Milano-Bicocca within a GlaxoSmithKline clinical research project BEL114054 (HGS1006-C1121). FMB is funded by the MEL-PLEX research training programme (‘Exploiting MELanoma disease comPLEXity to address European research training needs in translational cancer systems biology and cancer systems medicine’, Grant agreement no: 642295, MSCA-ITN-2014-ETN, Project Horizon 2020, in the framework of the MARIE SKŁODOWSKA-CURIE ACTIONS). DS was supported by the BioEntrepreneur-Fellowship of the University of Zurich (BIOEF-17-001).

## AUTHOR’S CONTRIBUTIONS

GC and MMB equally designed the experiments.

RT performed immunofluorescent tests, acquired the digital preparations.

MMB, MM and AMH devised the image analysis algorithms.

FMB and CP provided essential reagents and tissues.

FMB, MMB and MM performed visual and digital image analysis.

MF provided a customized version of the AMICO software and DS provided a pre-release version of histoCAT.

GC, AMH and MMB wrote the manuscript.

All authors have read and approved the final manuscript.

## METHODS

### MATERIALS AND METHODS

#### Tissues and Antigen Retrieval

Formalin fixed, paraffin embedded (FFPE) fully anonymous human leftover material used was exempt from the San Gerardo Institutional Review Board (IRB) approval as per Hospital regulations (ASG-DA-050 Donazione di materiale biologico a scopo di ricerca e/o sperimentazione, May 2012). Tonsil 1 and 3, belonging to two unrelated separate individuals, were processed in the very same fixation and embedding round. Tissue Microarrays (TMA) were constructed on a Galileo TMA CK4500 instrument (Integrated Systems Engineering, Milan, Italy) with 2mm cores of representative areas. Except when noted, a single 2mm TMA core was used for the analysis.

#### Immunofluorescent staining

Sections were processed for stainings essentially as described in (Bolognesi et al., 2017), with the following modifications:

- Sections were dewaxed using hexane (overnight) (Faolain et al., 2005) followed by xylene (10 min), rinsed in a graded alcohol series, rehydrated in distilled water (Scalia et al., 2017); antigen retrieval (AR) was performed as published (Bolognesi et al., 2017).
- Slides stained in IF were mounted with Phosphate buffered (pH 7.5) 60% Glycerol - 40% distilled water mixture containing 0.2% N-propyl Gallate and 584 mM sucrose (10% of a saturated solution), to which DAPI dilactate 5.45 μM (Sigma) were added; 10 mg of DAPI were dissolved in 2.19 ml of NN-dimethylformamide (Sigma) and the 10 μM stock solution was diluted to a 10 nM concentration in the PBS-glycerol. The concentration of DAPI was adjusted so that the DNA DAPI fluorescence did not bleed into the other channels.

Primary antibodies are listed in Table S1, secondary antibodies in Table S2. Primary antibodies were diluted at 1μg/ml in TrisHCl Buffered Saline (TBS) to which 2% BSA, 0.05% Sodium Azide and 100 mM Trehalose were added, after which sections were processed as per (Bolognesi et al., 2017).

#### Antibody stripping

Coverslips were gently removed by soaking the slides in TBS. Beta-mercaptoethanol / sodium dodecyl sulphate (2ME/SDS) stripping was performed as published (Bolognesi et al., 2017). After stripping, the slides were washed in TBS-Ts for at least 60 min with repeated buffer changes. No effect of staining sequence placement on exposure time or staining intensity was apparent, as previously published and shown in Figure S1G, H (Bolognesi et al., 2017). Complete removal of previous layers was monitored *a)* by checking for unexpected staining consistent with the subcellular and tissue distribution of a previously detected marker, *b)* by spurious co-clustering of unrelated molecules in the hierarchical clustering images (see below).

#### Image registration and digital subtraction of autofluorescence (AF)

Removing and replacing the very same slide on the scanner stage entails microscopic translations and rotations, which can misalign subsequently acquired images. To re-align subsequent images (register), one image of the nuclei in the tissue (DAPI) was set as the reference; then each other DAPI image acquired with a subsequent scan was aligned with the TurboReg Fiji plugin (Schindelin et al., 2012). The coordinates of the registration were recorded as landmark (a .txt file with coordinates) and applied to stacks of whole side images (sections or TMAs) with the Multi StackReg Plugin (Thevenaz et al., 1998). The process was automated with the A.M.I.C.O. software (Furia et al., 2013), which includes naming each image with biomarker’s name as detailed previously (Bolognesi et al., 2017)._AF was subtracted from FITC and TRITC channels as published (Bolognesi et al., 2017). AF subtracted images were used throughout for quantitative analysis.

#### Image Analysis

Representative manually selected areas, containing keratin+ surface epithelium, lymphoid follicles (composed of CD3+ T cells, PAX5+ B cells, etc.) and inner stromal core (Fig S1B), were sampled from tonsil WSI (average size 8.15 ± 1.65 sq mm) (Fig S1A) and from 2mm TMA cores (average size 3.14 sq mm). Grayscale 8 bit .tiff images were changed to 16 bit in batch with Fiji (Schindelin et al., 2012) before feeding into Cell Profiler (Carpenter et al., 2006) for mask creation and uploading to HistoCAT (Schapiro et al., 2017) for downstream image cytometry analysis. Preliminary analysis of lineage-specific stained, DAPI nuclear counterstained sections, revealed that in HLA-DR+ dendritic cells, Pax5+ large germinal center blasts and CD163+ macrophages the chromatin DAPI staining provided not enough contrast to serve as a reliable tool to identify these subsets by image threshold for segmentation (Figure S1C). This effect was evident also with a DAPI concentration 10x higher that what was currently used (not shown). We did not used a multimarker approach to cell segmentation (Schuffler et al., 2015) because *a)* our target population was supposed to be very small, thus requiring a very large number of segmented cells to be detected with this approach, *b)* which requires a nuclear marker and the low density of DAPI staining would not guarantee that cells with a complex profile would not be mistaken for empty spaces, *c)* our goal was to characterize S100AB+ cells. Therefore, the selected cellular markers, S100AB and HLA-DR, were employed for segmentation. Nevertheless, cell segmentation was attempted with the pan-leucocyte CD45 antigen, according to (Schuffler et al., 2015): DC clusters identified with the S100 and DR masks were identified, although with more noise, possibly because of the low DAPI content of some cells of interests (not shown).

An HLA-DR positive, CD20 negative mask (henceforth named DR-mask) was created by subtracting the CD20 image from the HLA-DR image with the Image Calculator function of Fiji (Schindelin et al., 2012). To ensure complete removal of the B cell marker, a 2.5x fold enhancement was applied beforehand to the CD20 image. A comprehensive S100AB mask (henceforth named S100-mask) was created on the S100AB image. First an image with the most intense cells was produced by threshold. Then the most intense cells from the image were removed with a threshold default algorithm and, after enhancing the resulting image by a factor ranging between 0.2 and 0.5 (depending on the image) both images were merged together in Fiji (Schindelin et al., 2012) and a mask obtained in Cell Profiler 2.2 (Carpenter et al., 2006). Both HLA-DR and S100 masks were created with CellProfiler and save with the .mat extension. Details of the Cell Profiler pipelines are reported in the Supplementary Information section. The output of the CellProfiler pipeline is an image composed of computationally segmented *n* objects, each one corresponding to a single segmented cell. The DR-mask identified an average 6566 cells in three tonsil areas, the S100-mask an average 9574.

The detection system used has a linear relationship between fluorescence output and antigen amount, differently from an amplified log ratio in flow cytometry. Intermediate-to-low levels of signal may tend to cluster in the negative area of the plot. Furthermore, we used an 8bit (256 channels) acquisition for time and storage space constraints. B cells are considered the reference population to set the gate for DC in the HLA-DR channel, and were optimally visualized in our samples. As a confirmation of the specificity and sensitivity of the assay, non-hematopoietic (see Figure S6D, E) and CD8+ T cell phenoclusters (not shown) were consistently HLA-DR negative and detected by the S100-mask only.

#### Bioinformatic processing

For each sample, all the images and one copy of either mask were placed in a dedicated folder and loaded into HistoCAT (Schapiro et al., 2017). Single cell mean intensity values, ranging from 0 to 255, for each marker were archsin transformed with cofactor 5 and then the values of intensity of all images were normalized with the dedicated function. For each tonsil, first tSNE dimensionality reduction technique (Amir et al., 2013; Maaten and Hinton, 2008; Schapiro et al., 2017) was used for low-dimensional visualization and afterwards Phenograph clustering (Levine et al., 2015) was applied to identify unique cell populations. Parameters were selected as suggested in the corresponding publications and implemented in HistoCAT. The analysis was performed with either mask independently by running a tSNE algorithm with all markers (minus DAPI), followed by running Phenograph (default parameter n=30). Single mask analysis and indirect evidence of HLA-DR and S100AB coexpression on discrete DC subsets, prompted an investigation with HistoCAT with both masks in separate folders, each containing the same set of images, followed by tSNE analysis of one folder vs the other, both selected.

A separated analysis was run with a mask created on CD123 image of whole tonsil #2 section (23616 x 13408 pixel) as well as on the same areas from the three tonsils examined with the S100- and DR-masks. Similarly, nuclear T-cell restricted TF LEF1 was used to create a T cell mask for every sample.

To analyze the sample with both S100 and DR masks, a Phenograph analysis was run first for each DR and S100 folders (n=30) as above and then with both folders together (n=30 and n=100) in order to better define cluster relationships among the two masks. Phenograph clustered cells obtained from single masks were superimposed on tSNE scatter plots, gated and visualized on the tissue images using HistoCAT and saved as separate images. To assign an immunologic and/or cell type identity to each cluster, four types of information were used: 1)The location along the X-Y coordinates in the Phenograph, absolute and relative to neighboring clusters, 2) The composite phenotype established by individual prototypic markers (e.g. CD1c, CD141,CD3, CD20, etc.) plus additional selected antigens. The granularity of the cluster composition was explored by hierarchical clustering within individual phenoclusters and a coloring of tSNE plots, 3) The composition in sub-groups identified by linear hierarchical clustering within each cluster, and 4) The distribution on the tissue, with respect to anatomically and phenotypically distinct microenvironmental areas (stromal axis, T cell, B cell, subepithelial, intraepithelial areas, germinal centers, vessels, etc.).

Phenogroups with a selective tissue distribution in areas occupied by previously identified DC types were verified for engagement by plotting each group on two-color images and by co-plotting on tissue images T-cell and DC phenogroups, these obtained with a combined LEF1, S100 and DR masks.

The reproducibility between samples of the same tissue was tested 1) by uploading two 2 mm core replicas for each of 10 reactive lymphoid tissue and running a tSNE analysis with the DR-mask, 2) analyzing a second set of tonsil samples on TMA cores and comparing the cell type clusters. Results from the first approach show that duplicates overlap (not shown).

#### Heatmaps in R

We made heatmaps using R version 3.3.3 (R Core Team (2017). R: A language and environment for statistical computing. R Foundation for Statistical Computing, Vienna, Austria. URL https://www.R-project.org/). We built an R script to process the histoCAT output data (available in the Supplementary Information section). At first, we set up an R working directory that included the two comma separated values (.csv) files of the S100 and the DR-masks, which contained raw values and annotations regarding phenograph clusters for each cell. The .csv files were imported in two dataframes, subsequently joined by raw binding, and furtherly processed to obtain curated data. The dataframe was transformed in a matrix of values, and a default standardization (centering and scaling of values) was performed by the *scale* function of base R package #1 (ibid.). Heatmaps were made using the *ComplexHeatmap* #2 package (Gu et al., 2016) so that each phenograph cluster was separately visualized and annotated. All cells within each phenograph group were hierarchical clustered by a dendrogram which used the Ward’s D2 algorithm. All column markers were organized in a hierarchical way through the same algorithm. An additional color-coded column allowed a clear visualization of the mask affiliation for each heatmap phenograph cluster.

#### Quantitation of DC subsets

The number of computationally segmented cells in each phenocluster for each mask was the numerator. In case of totally or overlapping masks, the higher cell number was used, the sum in any other case. Two denominators were used: the total number of segmented cells in each mask or the total number of nuclei identified by DAPI thresholding in the analyzed areas, to which the total segmented cell number was added, this latter because of the inability to identify DCs by DAPI nuclear mask (Figure S1C).

#### Fluorescent tissue images and localization of phenoclusters in tissue

Stacks containing all the images were cropped for regions of interest (ROI) representing the cell type sought and by the tissue location of the phenocluster provided by HistoCAT. The single stain greyscale images from the stack were inverted, optimized in ImageJ by an enhancing factor comprised between 0.3% and 0.01% and overlayed with a two-color rendition of the original masks. The tissue distribution of individual phenoclusters produced by HistoCAT as jpg files was extracted by wand color-selection in Adobe Photoshop CS3 (Adobe Systems Incorporated, San Jose, CA), coded with the mask color (S100 or HLA-DR) and superimposed on whole area images composed of immunofluorescent CD4, CD8 and DAPI in inverted, subdued colors.

## SUPPLEMENTARY INFORMATION

The landscape of S100B+ and HLA-DR+ dendritic cell subsets in tonsils at the single cell level via high-parameter mapping.

Maddalena Maria Bolognesi, Francesca Maria Bosisio, Marco Manzoni, Denis Schapiro, Riccardo Tagliabue, Mario Faretta, Carlo Parravicini, Ann M Haberman, Giorgio Cattoretti

**TABLE S1.**
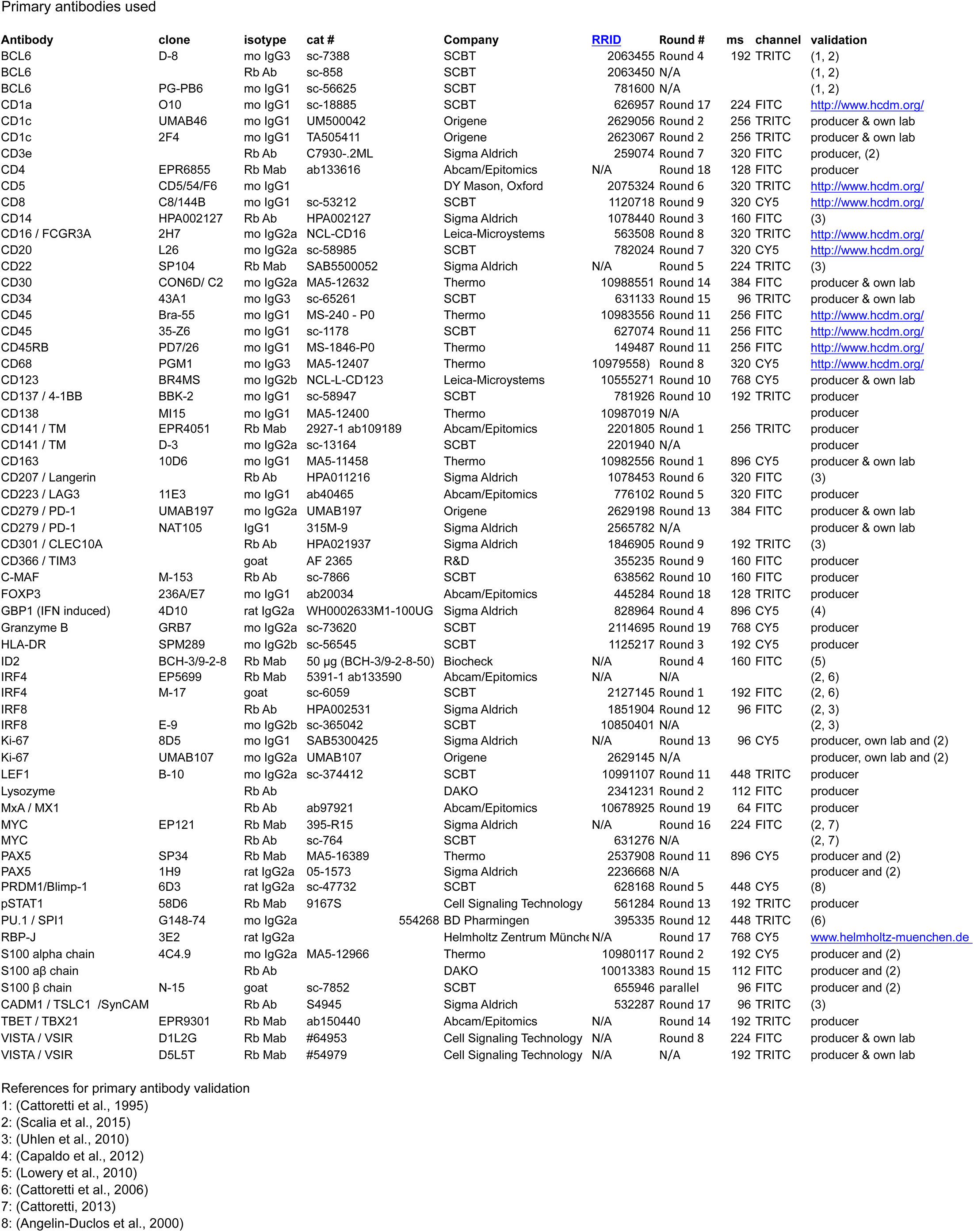
Primary antibodies used

**TABLE S2.**
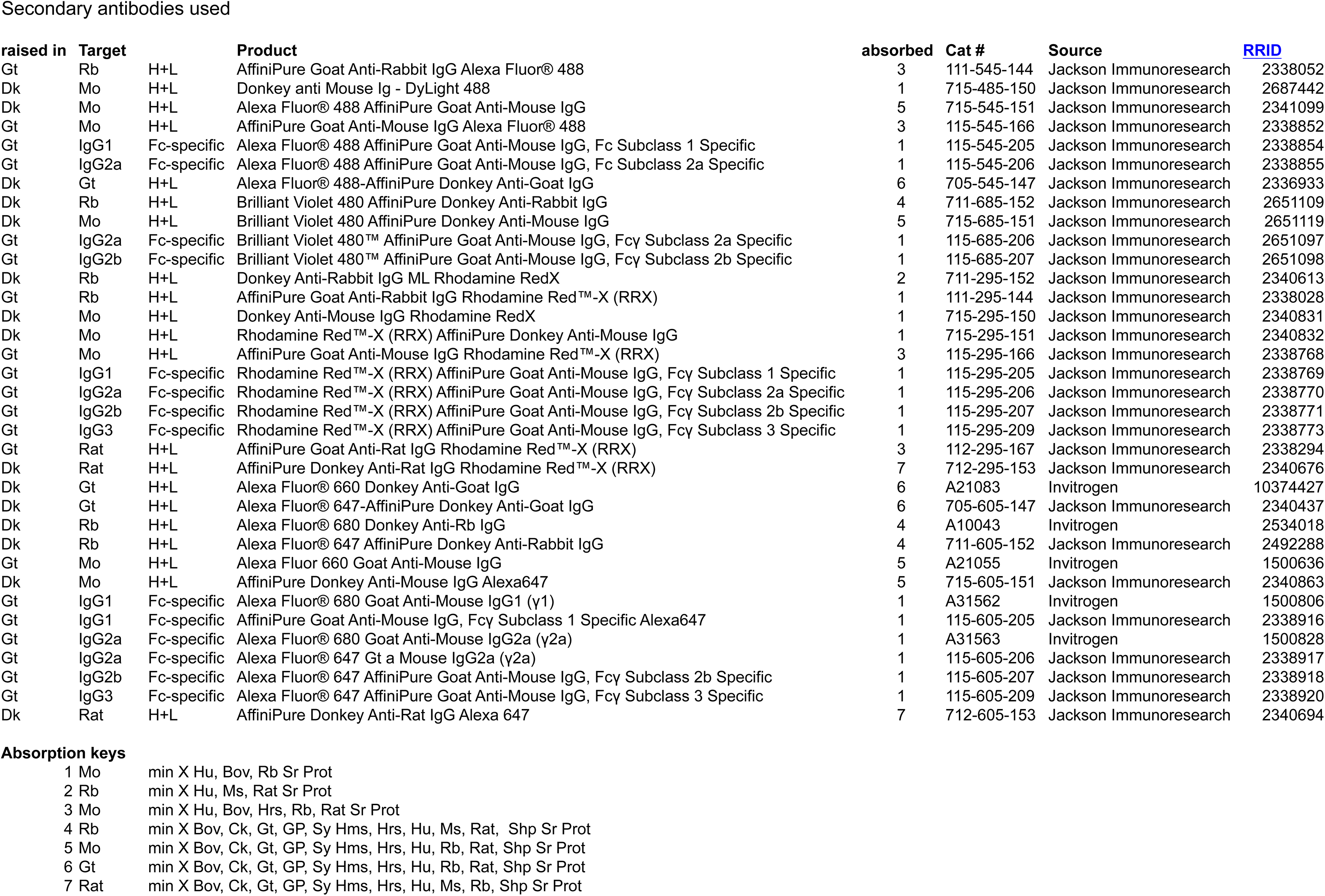
Secondary antibodies used

**Supplementary Table 3.**
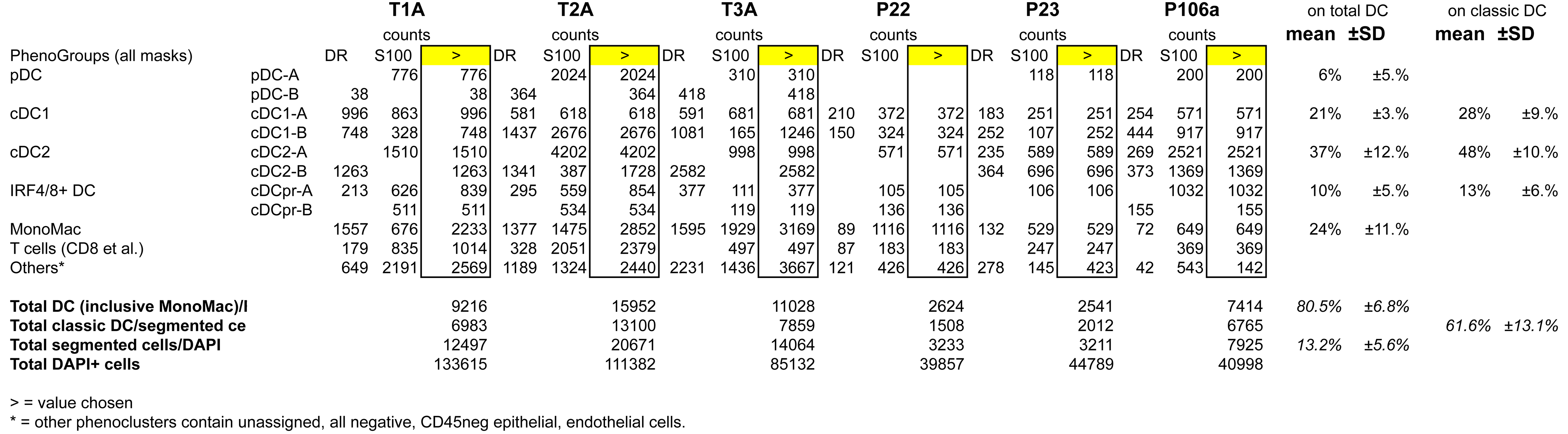

**Figure S1.**
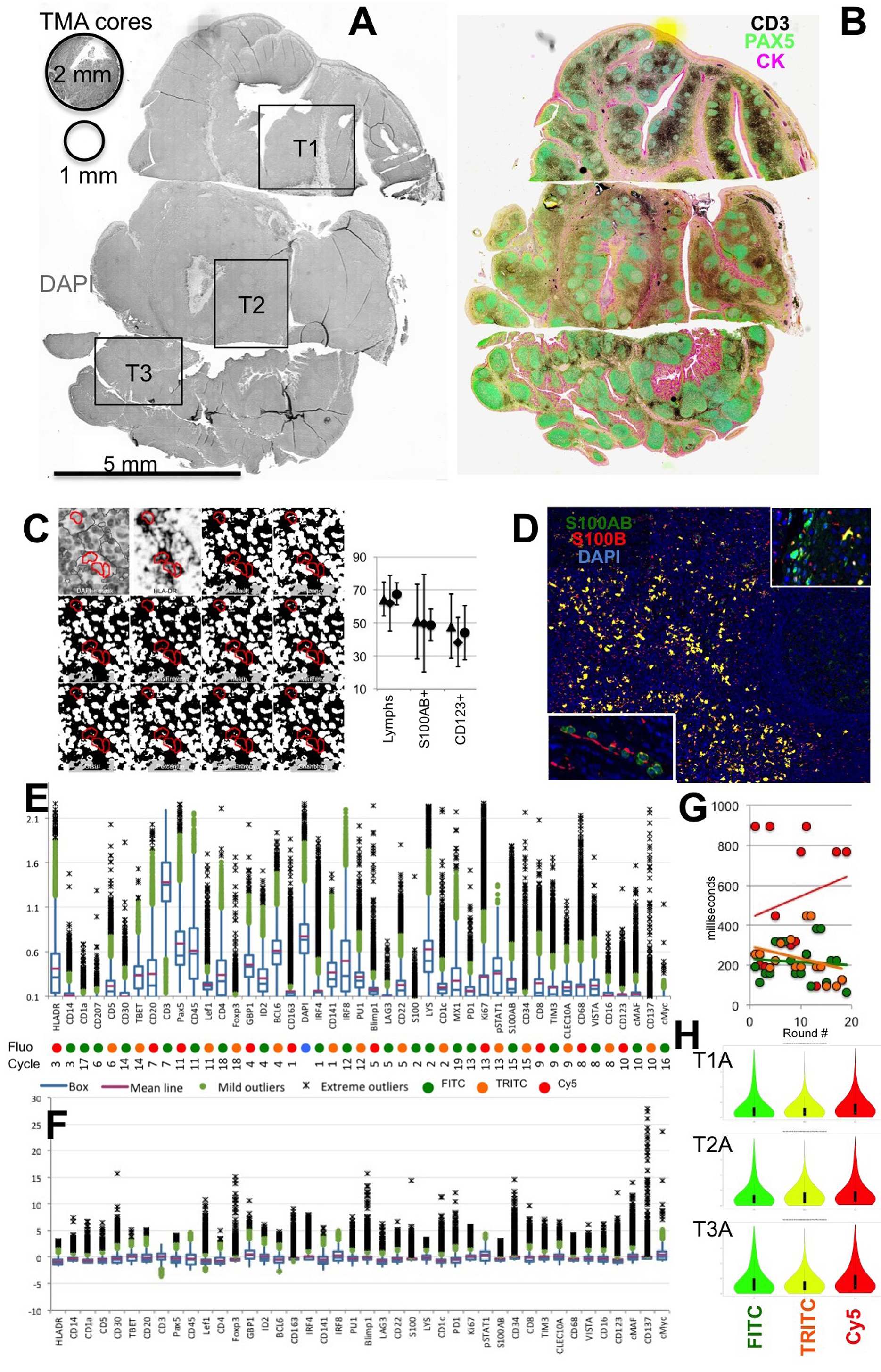
Landscape of the tissue sections, thresholding and cell identification, quantitative fluorescence parameters. Related to Fig.1. **A**: Low power image of the inverted whole slide DAPI stain of the three tonsil sections. Each square indicate the area acquired and the tonsil name. On the top left, the size of a 1 and 2 mm TMA cores are depicted. **B**: a serial section to A is stained for T cells (CD3, black), B cells (PAX5, green), Cytokeratins (pink) and DAPI (yellow) and shown at low power as an inverted color composite. **C**: A representative 73×76 μm field shows an inverted DAPI image, a positive HLA-DR stain on DC, a DR-mask (red) superimposed to 10 different thresholding algorithm for DAPI (additional 5 algorithms not shown). Note the faint DAPI nuclear stain and the absence of DAPI mask inside the HLA-DR+ DC. The mean ± SD DAPI intensity on a 0-255 gray levels scale is shown for all lymphocytes (Lymphs), S100AB+ dendritic cells and CD123+ plasmacytoid DC (pDC). CD8+ T lymphocytes were not removed from the S100 mask used for DAPI measurement, hence the larger standard deviation. **D**: A low power image of a tonsil area sequentially stained for rabbit anti S100AB (green), goat anti S100B (red) and DAPI (blue) nuclear counterstain. The vast majority of cell coexpress S100AB and −B (yellow). In the insets, S100A1+ myeloid cells (bottom) and terminally differentiated squamous epithelial cells (top). **E**: Box plot of the arcsin transformed (HistoCAT) fluorescence values for each stain for all the 8222 segmented cells of the S100-mask in tonsil 1. The fluorochrome and the cycle position of each marker is shown below each marker. **F**: Box plot of the normalized data of the fluorescence values for each stain as per **E**. **G**: Exposure time (in milliseconds) vs staining cycle position for each marker, according to the fluorescence channel, color-coded as per **E**. The intercept for each fluorescence is plotted. **H**: violin plots of the normalized fluorescence values for each fluorescence channel and for each of the three tonsils.

**Figure S2.**
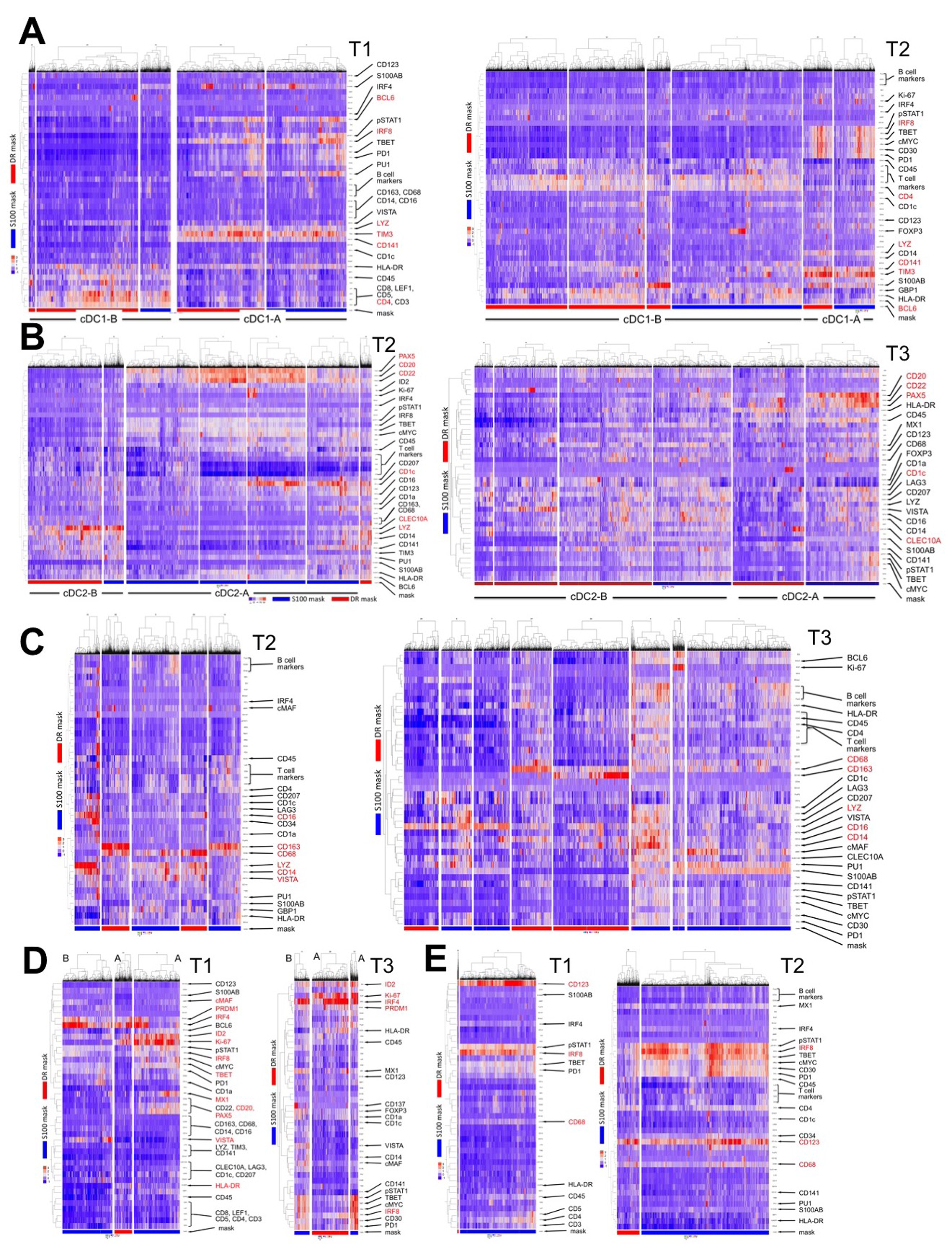
Extended heatmap profiles of DC Phenoclusters Related to Fig. 2, 3, 4, 5 and 6. The image contains Phenocluster-specific heatmap profiles for samples not shown in Fig. 1-5. Phenograph clusters from the Tonsils not shown in Figures 1-4 are shown. Each phenocluster is hierarchically clustered by markers (rows) and individual segmented cells (colums); the mask by which each phenocluster has been obtained is exemplified by the color band at the bottom: S100AB: blue, HLA-DR red. Markers with arrows listed on the right are DC lineage and positive markers highlighted by the dendrograms. Distinctive and characterizing markers are in red. **A**: cDC1; **B**: cDC2; **C**: MonoMac; **D**: IRF4/8+ DC; **E**: pDC

**Figure S3.**
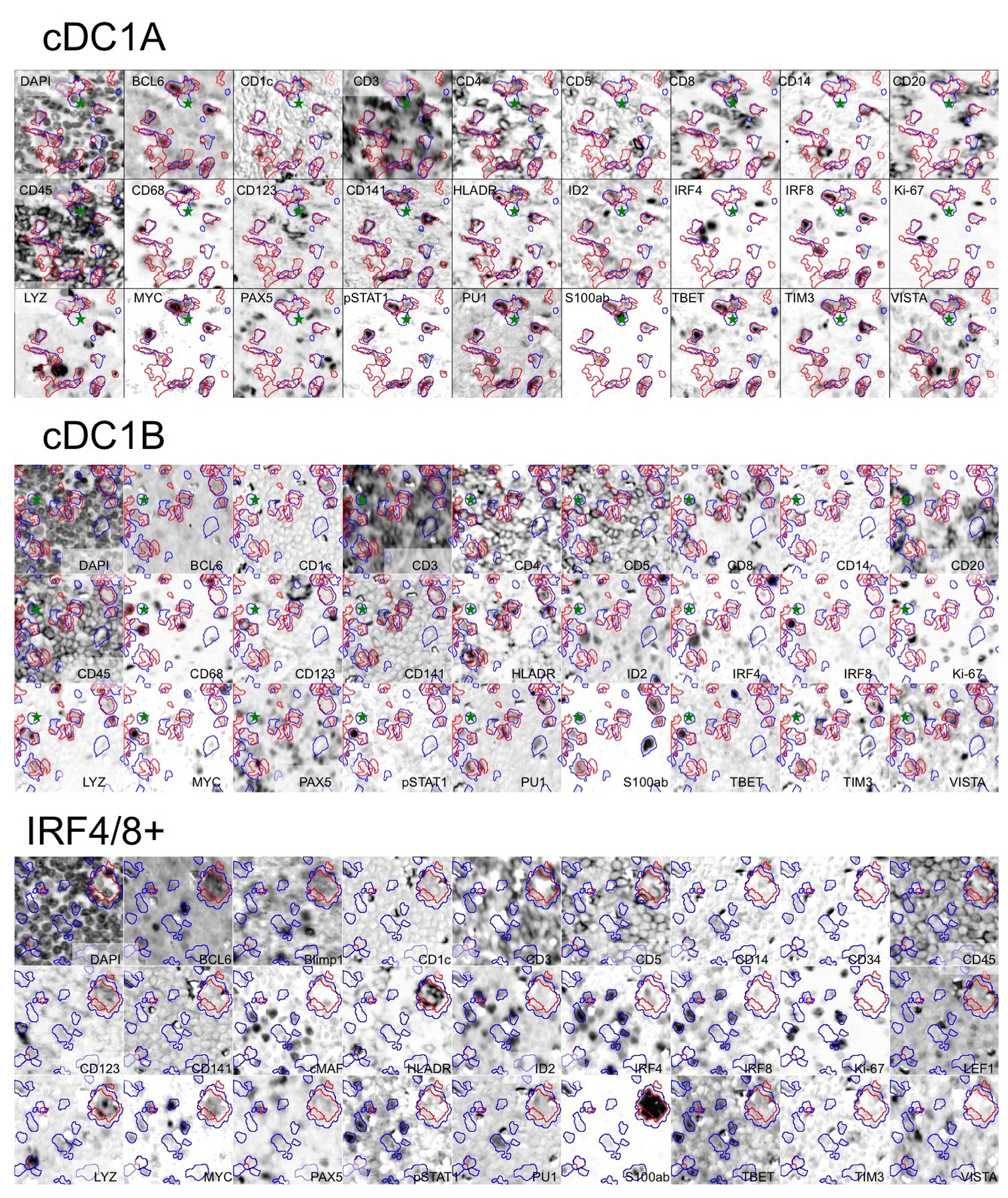
DAPI and 28 marker high-power images for cDC1 and IRF4/8+ dendritic cells Related to Fig. 2 and 4. Images of selected stains, representative of the cDC1-A, cDC1-B and IRF4/8+ DC population, are shown as inverted fluorescent images; the color-coded outline of the S100AB and HLA-DR mask are superimposed. The stain name is written below each image. In cDC1-A and −B an asterisk marks CD8+ S100+ T cells. Images size 89×89 μm for cDC1-A and −B, 70×70 μm for IRF4/8+ DC.

**Figure S4.**
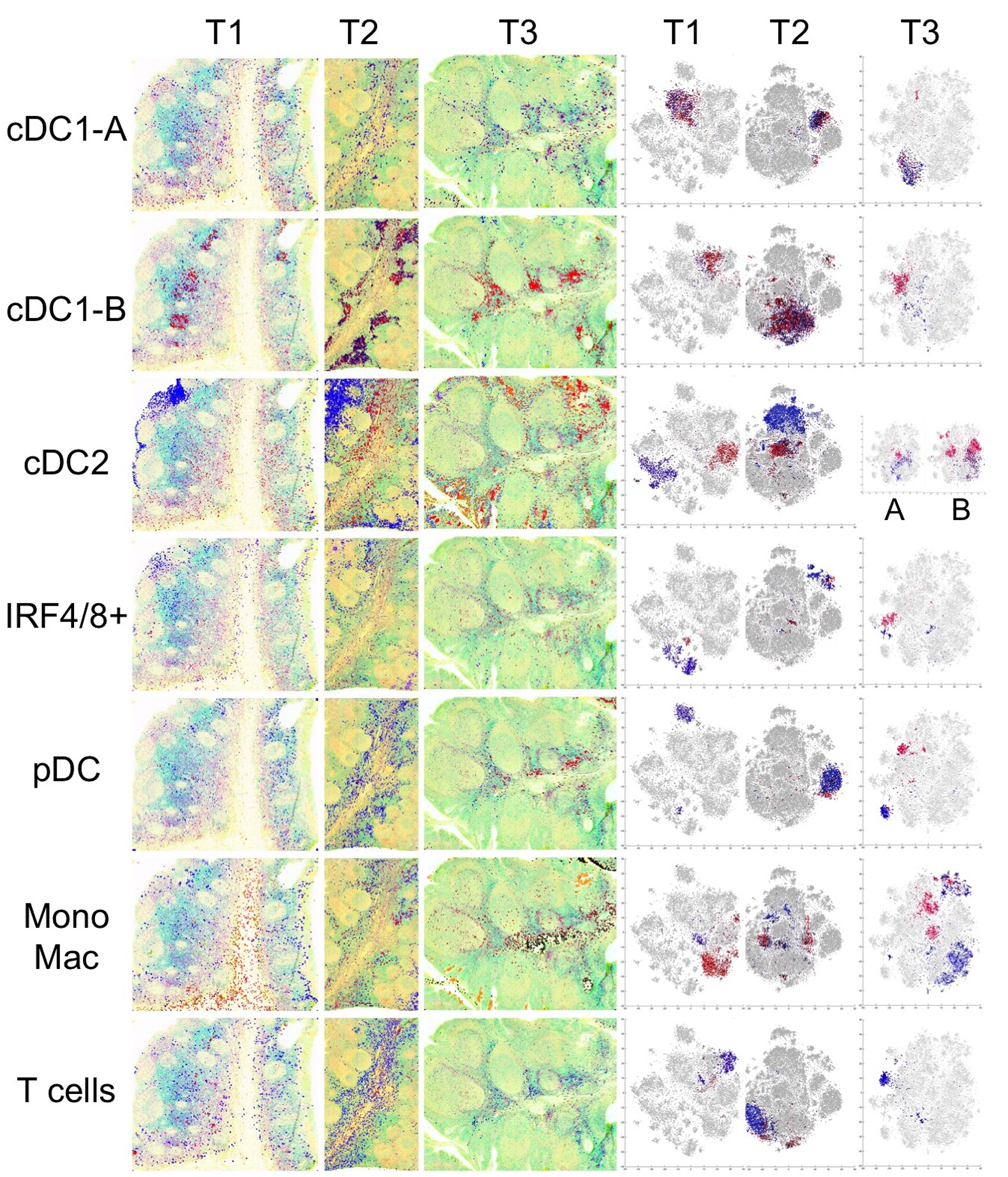
Tissue distribution maps of the DC phenoclusters and allocation in the tSNE silhouette. Related to Fig. 2, 3, 4, 5 and 6. LEFT panel: Tissue distribution of Phenograph clusters from the Tonsils not shown in Figures 1-4 are shown. The tissue image is rendered by a tenuous CD4, CD8 and DAPI staining. Each population is plotted with a color matching the mask from which the data are derived. T cells are S100AB+, largely CD8+ T cells. RIGHT panel: The distribution of each phenocluster for the Tonsils not shown in Figures 1-4 are shown. Each cluster is highlighted with the dot color matching the mask color. For clarity, cDC2 phenoclusters are shown separately for cDC2-A and −B

**Figure S5.**
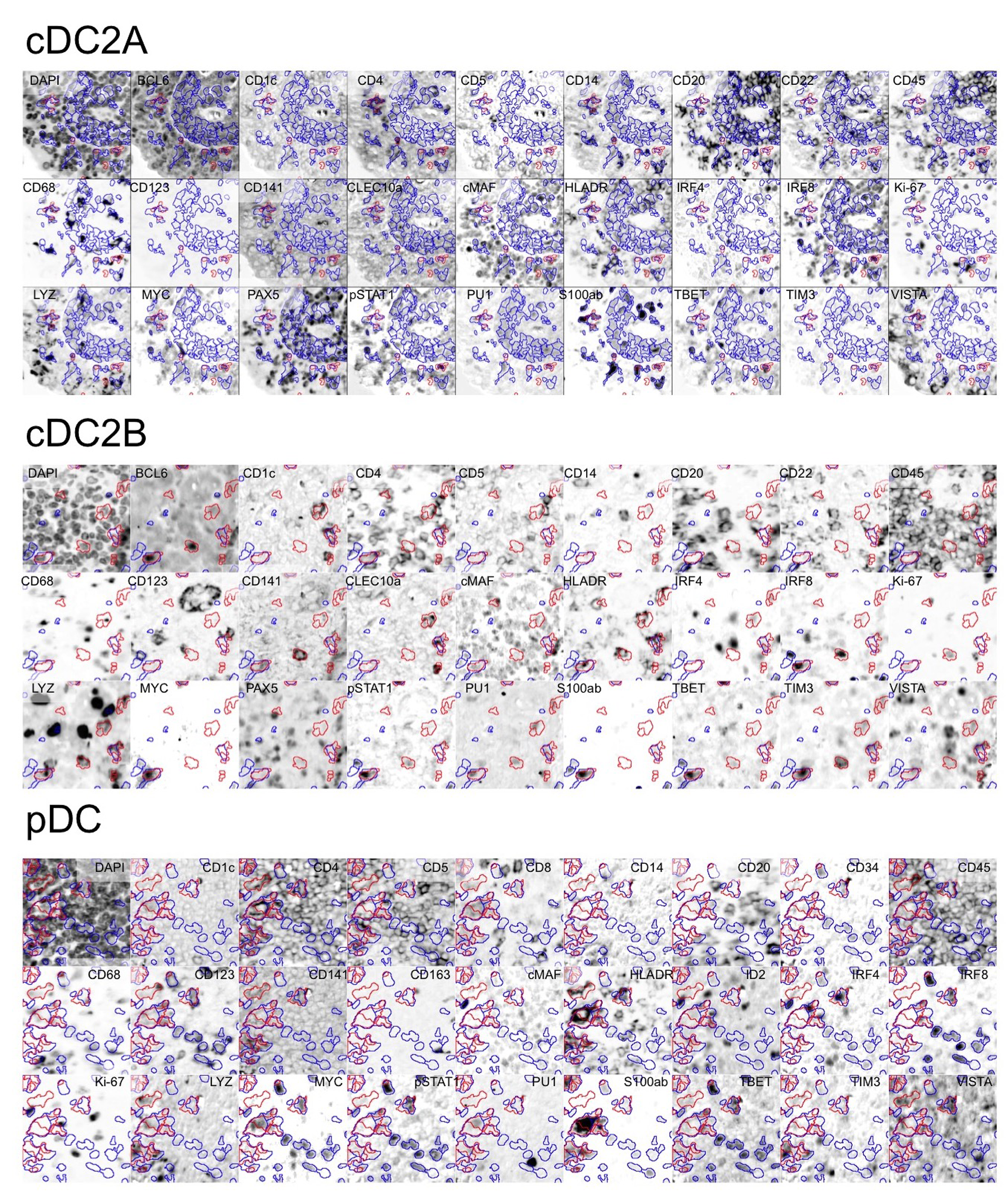
DAPI and 28 marker high-power images for cDC2 and pDC dendritic cells Related to Fig. 3 and 5. Images of selected stains, representative of the cDC2-A, cDC2-B and pDC population, are shown as inverted fluorescent images; the color-coded outline of the S100AB an DR masks are superimposed. The stain name is written below each image. Images size 118×118 μm for cDC2-A, 89×89 μm for cDC2-B, 84×84 μm for pDC.

**Figure S6.**
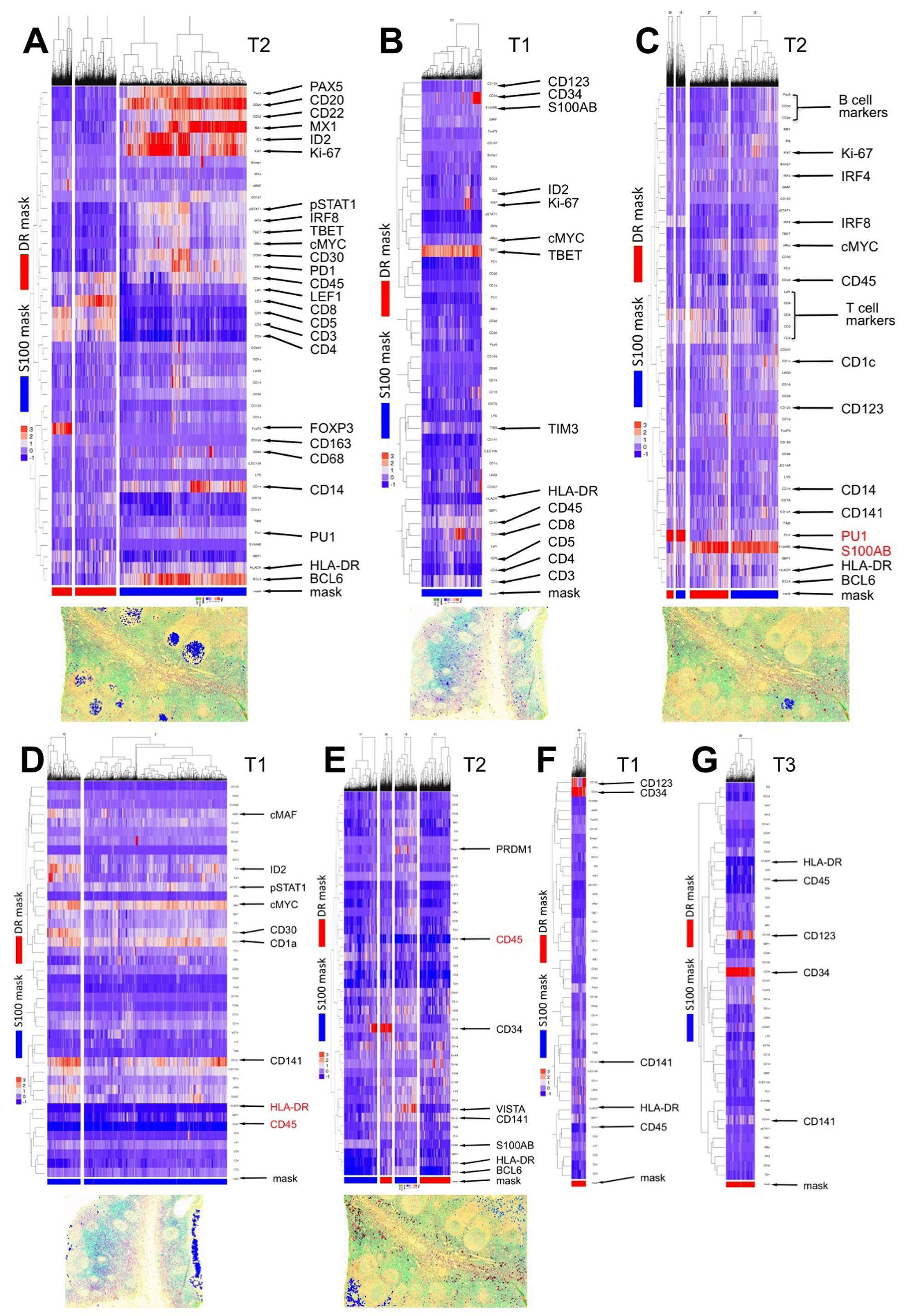
Detailed phenotype and tissue localization for minor cell subsets identified in isolated tonsils. Related to Fig. 1. **A**: RIGHT: a Germinal Center B cell subset (B cell markers+, Ki-67+, BCL6+) is identified by neighboring CD14+ follicular dendritic cells and an S100-mask. The tissue distribution is shown below. LEFT: HLA-DR+ CD8+ and FOXP3+ CD4+ T cells are identified by the DR-mask. **B**: a TBET+ population devoid of other DC markers except S100AB is seen in Tonsil 1. The tissue distribution is shown below. **C**: PU1+ and S100AB strongly positive cells adjacent to T cells are identified in Tonsil 2. No other defining marker is present. **D**: HLA-DR and CD45 negative epithelial cells are identified in Tonsil 1. This group is distributed at the crypt surface. **E, F, G**: CD45 negative CD34+, CD123±, CD141± endothelial cells are identified in Tonsils 1, 2 and 3.

### SUPPLEMENTARY DATA

Cell Profiler files

R script Heatmap

.csv files

Go to:
https://data.mendeley.com/

